# Identification of RNF114 as ADPr-Ub reader through non-hydrolysable ubiquitinated ADP-ribose

**DOI:** 10.1101/2025.05.08.652854

**Authors:** Max S. Kloet, Chatrin Chatrin, Rishov Mukhopadhyay, Bianca D. M. van Tol, Rebecca Smith, Sarah A. Rotman, Rayman T. N. Tjokrodirijo, Kang Zhu, Andrii Gorelik, Peter A. van Veelen, Dragana Ahel, Ivan Ahel, Gerbrand J. van der Heden van Noort

## Abstract

Crosstalk between the post-translational modification processes ubiquitination and ADP-ribosylation occurs in DNA-damage and immune-responses, in addition the physical linkage of ADP-ribose and ubiquitin is found during bacterial infection. Here, we study the ubiquitination of ADP-ribose mediated by human Deltex E3 ligases and the subsequent fate of the formed hybrid post-translational modification. We prepare a non-hydrolysable ADPr-Ub probe that we employ in a chemoproteomics approach and identify RNF114 as an interacting protein. Using biophysical and biochemical experiments, we validate that RNF114 preferentially interacts with ubiquitinated ADP-ribose over non-modified ubiquitin. Subsequently, RNF114 can elongate the ubiquitinated ADP-ribose with a K11-linked ubiquitin chain. Using domain deletion analysis, we pinpoint the tandem zinc fingers and ubiquitin interacting motif (ZnF2+ZnF3+UIM) domains of RNF114 to be crucial for recognising ubiquitinated ADP-ribose. Moreover, these domains are essential for the recruitment of RNF114 to the sites of laser-induced DNA damage.

## Introduction

Ubiquitination is a much-studied post-translational modification (PTM) playing crucial roles in regulating cellular processes across eukaryotes. Canonical ubiquitination relies on a cascade of E1-, E2-, and E3 enzymes which conjugate the Gly76 C-terminus of ubiquitin (Ub) to the lysine side chains in proteins creating a stable isopeptide bond(*1*). However, it became evident that Ub can also be conjugated to non-lysine residues in proteins to form a thioester bond with cysteine(*2, 3*) or an oxyester bond with serine or threonine(*4–6*). Beyond protein modification, recent discoveries reveal ubiquitination on several non-protein substrates, such as lipopolysaccharide(*7*), oligosaccharide(*8*), phosphatidylethanolamine(*9*), nucleic acids(*10, 11*), as well as on another PTM; ADP-ribose (ADPr) (*12, 13*).

ADP-ribosylation is an ancient modification which is conserved across all kingdoms of life. This modification relies on ADP-ribosyltransferase enzymes which utilise NAD^+^ and transfer ADP-ribose onto proteins and nucleic acids(*14, 15*). Crosstalk between ubiquitination and ADP-ribosylation occurs in mammals where for instance poly-ADP-ribose (PAR) chains direct subsequent ubiquitination events in DNA damage responses(*16*), Wnt signaling (*17*), and ADPr-dependent ubiquitination in immunity responses (*18*). Such crosstalk is also cleverly utilised by bacteria, which themselves lack Ub, to disrupt host Ub signaling by modifying it with ADPr, as shown by *Legionella* and *Chromobacterium*(*19, 20*). These are prime examples where the already complex ubiquitination code is made even more diverse upon crosstalk with ADPr.

In humans, the family of Deltex RING E3 Ub ligases (DTX1, DTX2, DTX3, DTX3L, DTX4) catalyse Ub conjugation to the 3’-hydroxyl group of ADPr, which results in a hybrid ubiquitinated ADP-ribose modification (further annotated as ADPr-Ub) (**Fig. 1A**)(*12, 13, 21*). It was first reported that the heterodimeric complex of DTX3L/PARP9 conjugates ADPr to the Gly76 residue of Ub(*22*). This unique reaction is catalysed by all members of Deltex E3 ligases through their conserved RING and DTC domains, which interacts with Ub-loaded E2 (E2∼Ub) and NAD^+^/ADPr, respectively(*21*). The DTC domains of Deltex E3 ligases bind to NAD^+^ and ADPr, and can recruit ADP-ribosylated proteins for ubiquitination(*23*). However, instead of ADP-ribosylation of Ub which occurs on the C1” of nicotinamide ribose, Deltex E3 ligases catalyse the ubiquitination of ADPr on the 3’-hydroxyl of adenosine ribose (Fig 1A) (*12*). To support these reactions, the DTC domains of Deltex E3 ligases harbour a conserved Glu-His diade to deprotonate the 3’-hydroxyl of the proximal adenosine ribose in ADPr to attack the thioester E2∼Ub conjugate, resulting in a covalent ester bond between Ub Gly76 and the ADPr 3’-hydroxyl group(*12*). This allows formation of dual hybrid modification composed of ubiquitin and ADPr (ADPr-Ub) either on protein or nucleic acids substrates(*12, 13*). However, the biological function of such Deltex-catalysed hybrid ADPr-Ub-substrates remains unknown.

**Figure 1.**
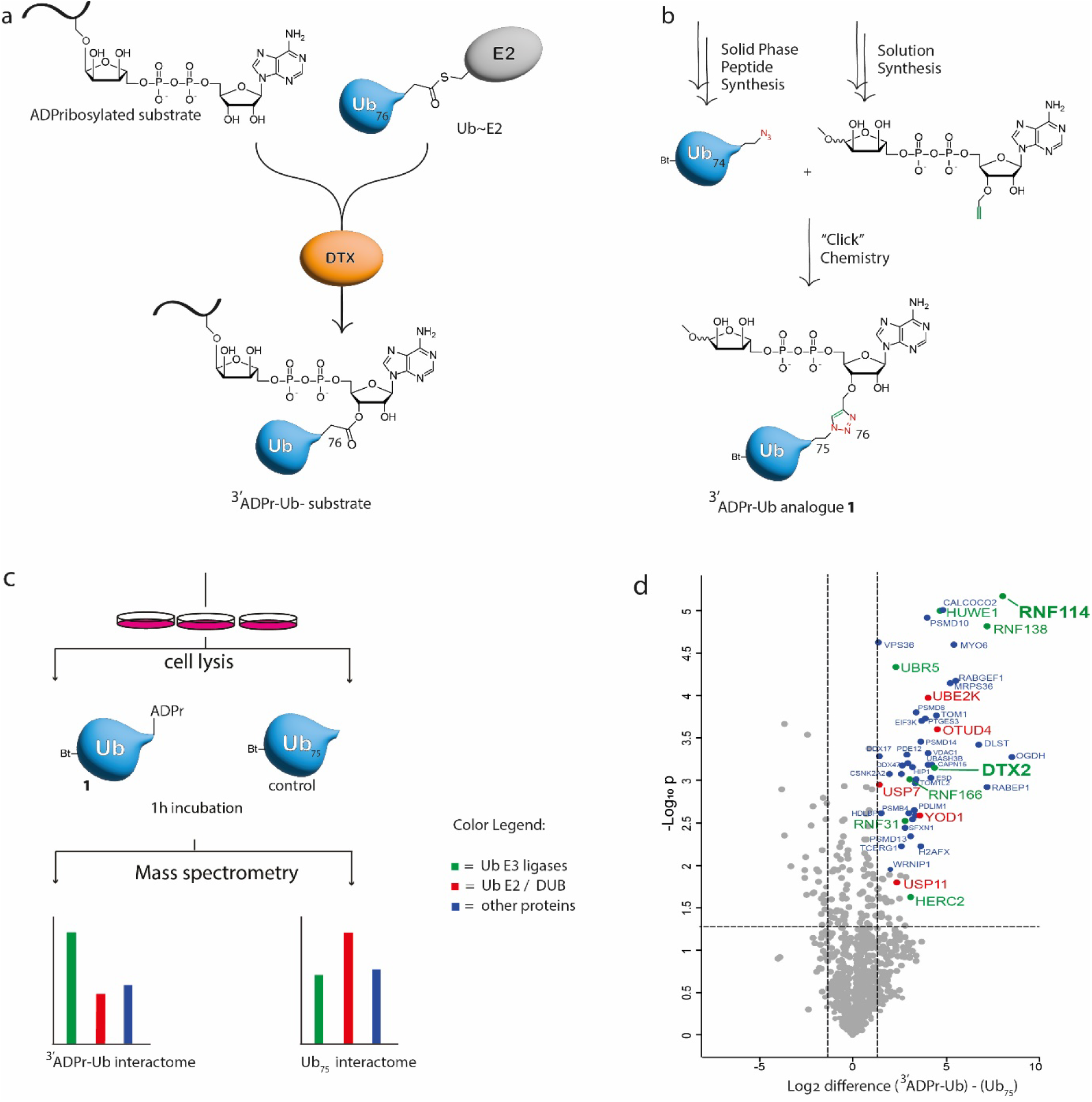
Quantitative chemo-proteomics identifies interactors of ADPr-Ub probe 1. **A)** Schematic representation of the conjugation activity of DTX enzymes. **B)** The chemical synthesis scheme to obtain **1** via CuAAC between a 3’-*O* propargylated ADPr and Ub_1-74_ bearing a C-terminal azide. **C)** Schematic representation of the workflow for quantitative chemo-proteomics. **D)** Volcano plot showing the interactome of probe **1**. RNF114 and DTX2 are highlighted in **bold**. Data is normalized to the Ub_75_ interactome.

In this work, we wondered if there are proteins involved in reading this new expansion of the ubiquitin code, which would establish a first indication regarding the biological consequence and the underlying molecular mechanisms of the hybrid ADPr-Ub. One of the major challenges in studying the ADPr-Ub modification is the inherent lability of the ester bond. Here we synthesised a non-hydrolysable probe that mimics the formed ubiquitinated ADP-ribose to perform an unbiased pulldown from cell lysate and start exploring its interactome. We identified RNF114 as one of the top binders for ADPr-Ub. RNF114 was previously suggested to bind mono-ADP-ribose (MAR) and poly-ADP-ribose (PAR) through its Zinc finger (ZnF) domains (*24, 25*). We discovered that RNF114 mediates low micromolar binding to ADPr-Ub through its tandem ZnF2+ZnF3 and Ub-interacting motif (UIM) domains. We further showed that RNF114 can engage ADPr-Ub and elongate it with K11-linked polyubiquitin chains forming a novel mixed Ub chain linkage product. Finally, the tandem ZnF2+ZnF3+UIM domains of RNF114 are indispensable for its recruitment to the sites of DNA damage in cells, which suggests RNF114 as a reader of ADPr-Ub signal during response to DNA damage.

## Results

### Synthesis of a ADPr-Ub analogue

The activity of Deltex enzymes links the C-terminal Gly76 carboxylate of Ub to the 3’ proximal hydroxyl group of adenosine in ADPr or NAD^+^, forming an ester bond. ADPr-esters have been shown to be chemically labile (*26*) and prone to acyl migration (*27*). To prevent this and unwanted (enzymatic) hydrolysis when probing for interactors in cell lysate we envisioned a triazole linkage between ADPr and Ub to be the ideal stable linkage. Additionally, copper-catalyzed azide alkyne cycloaddition (CuAAC) reactions between propargylated ADPr and a Ub-mutant equipped with an azide were shown to be an effective method to install the ADPr moiety on Ub as the final step in synthesis, thereby reducing either acid or alkaline induced problems during chemistry (*28*). Considering these advantages, the synthesis of a ADPr-Ub analogue **1** was undertaken, establishing an inherently stable triazole linkage between Ub and ADPr (**Fig. 1B).** We used solid phase peptide synthesis to prepare the Ub protein and decorated the N-terminus with biotin to facilitate pulldown experiments and immobilization of the probe. After solid phase peptide synthesis, we installed an azido group via a short linker on the C-terminal tail of Ub. We introduced an alkyne on the 3’-alcohol of 1”-OMe-ADPr, a proxy of an ADP-ribosylated serine residue, by preparing a 3’-O-propargylated adenosine amidite that was conjugated to a suitably protected ribose-5”-O-phosphate via P(III)-P(V) methodology (*29*) (**Supplementary** Fig. 1). To facilitate the complete in-solution deprotection by aqueous ammonia, both fragments were protected with base-labile protecting groups.

### Identification of ADPr-Ub readers using chemo-proteomics

In order to validate whether ADPr-Ub probe **1** can be used to profile the interactome of ubiquitinated ADPr we performed a pulldown from HEK293T cell lysate and immunoblotted against DTX2 (**Supplementary** Fig. 2). We confirmed DTX2 was enriched using probe **1** whereas the negative control Ub_75_ was not able to enrich DTX2. Ub_75_ lacks both the ADPr moiety and the most C-terminal Gly76 residue preventing it from being conjugated to substrates via the normal conjugation machinery. To identify relevant ADPr-Ub interactors in cells, a pulldown experiment using probe **1** was performed including Ub_75_ as negative control (**Fig. 1C**). Cells were lysed using CO-IP buffer and incubated with either probe or control followed by biotin pulldown using Neutravidin beads. Pulldown samples were washed with CO-IP buffer under mild conditions and subjected to LC-MS/MS analysis. We normalized the data against proteins interacting with the Ub_75_ control and indeed also confirmed the validity of this approach by identifying DTX2. Proteins most significantly interacting with probe **1** include RNF114 and RNF138, both Ub E3 ligases have been linked to DNA damage repair pathway, regulation of ADP-ribosylation events, and immunity responses (*24, 25, 30–32*) (**Fig. 1D**). Other interactors include OTUD4, USP7, YOD1 and USP11 reported as “erasers” of ubiquitination while interactors like UBR5, HUWE1, UBE2K, HERC2, RNF166 and RNF31 are reported as “writers” of ubiquitination. These proteins all exhibited a ≥ 1.3-fold difference of the Log_2_ intensities and Log_10_ P value significance. Some of these hits (DTX2 and HUWE1) contain a WWE domain, which binds to the iso-ADPr moiety of PAR (*33*). To demonstrate global variation among the three replicate samples and detect unknown patterns across various conditions, we employed principal component analysis (PCA). The PCA profile revealed clustering of replicates from different sample groups (**Supplementary** Fig. 3A). Furthermore, hierarchical clustering analysis demonstrated different protein interaction profiles for both probes (**Supplementary** Fig. 3B), where we highlighted the interaction profile of DTX2 and RNF114 (**Supplementary** Fig. 3C,D). Analysis of the ADPr-Ub interactome revealed multiple unique protein interactors of ADPr-Ub including RNF114, RNF138, RNF166 and DTX2.

### RNF114 and DTX2 RING-DTC (RD) bind ADPr-Ub

We next investigated RNF114 as the most significant interactor from the proteomics data to verify bona-fide binding to biotin-ADPr-Ub (probe **1**) using immunoblotting. We also investigated DTX2, which was identified in our dataset and is one of the writers of ADPr-Ub modification. For this experiment, probe **1** and the negative control (biotin-Ub_75_) were incubated with cell lysate. A pulldown with Neutravidin beads followed by immunoblotting indeed enriched both RNF114 as well as DTX2 by probe **1** and not by the Ub_75_ control (**Fig. 2A**). Next, we sought to further validate the interaction between probe **1** and RNF114 and used bio-layer interferometry to determine their binding affinities. RNF114 as well as RNF138 and RNF166, that come forward as significant binders to probe **1** in our proteomic data have been identified previously as binders of ADPr transferase Tankyrase 1(*32*). Both RNF114 and DTX2 show no detectable binding to Ub or mono-ADPr, but show a strong interaction with ADPr-Ub (**Fig. 2B** and **Supplementary** Fig. 4). Using a titration range, we were able to deduce apparent K_D_’s of 850 nM for RNF114 and 1.4 µM for DTX2 (**Fig. 2C**).

**Figure 2.**
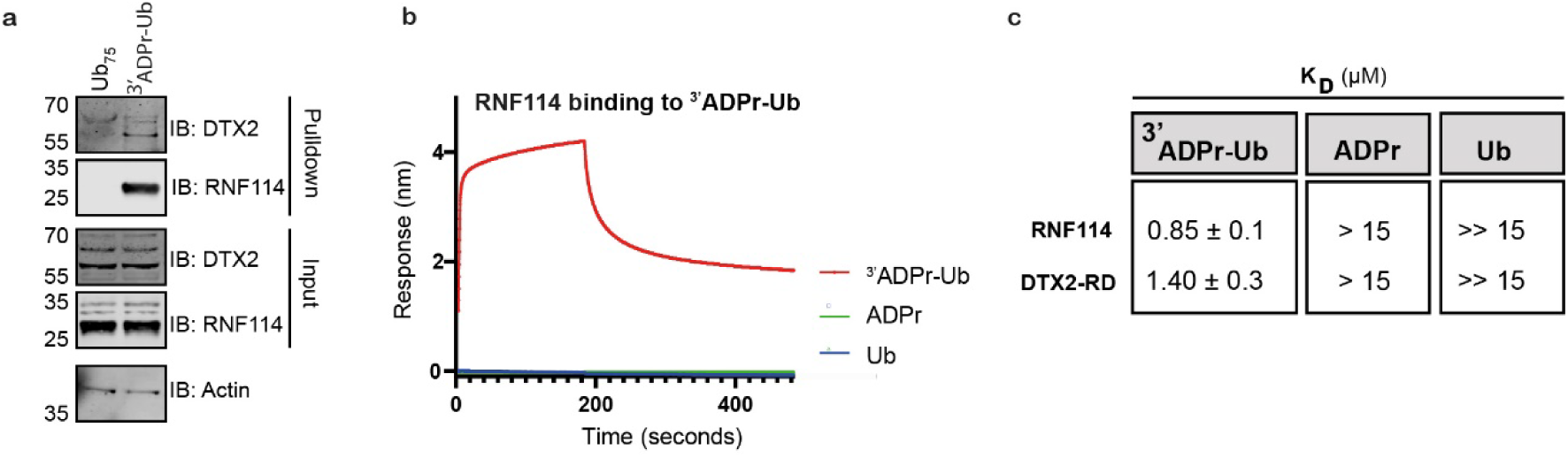
DTX2 and RNF114 interacts with ADPr-Ub. **A)** Both biotin-Ub_75_ and biotin-ADPr-Ub probe **1** were incubated in cell lysates from HEK293T cells then pulled down using Neutravidin beads. Western blot shows levels of endogenous DTX2 and RNF114 interacting with the probes where Actin serves as the loading control, **B)** Bio-layer interferometry response curves of RNF114 to biotin-Ub_75_, biotin-ADPr-Ub or biotin-ADPr, **C)** Apparent K_D_ binding affinities of RNF114 and DTX2 deduced from BLI measurements.

### RNF114 elongates ADPr-Ub

Our results thus far suggest that RNF114 and DTX2-RD selectively bind ADPr-Ub over Ub or ADPr in isolation. While the RD domains of Deltex proteins, including DTX2, are able to catalyse Ub transfer to ADPr or NAD^+^ in the presence of ubiquitination cascade components, the role of RNF114 in relation to ADPr-Ub is unclear. Unlike DTX2, RNF114 is unable to catalyse such adduct formation when tested with ^32^P-NAD^+^ and ubiquitin cascade components (**Fig. 3A, Supplementary** Fig. 5A). To our surprise, when both DTX2-RD and RNF114 are present in the reaction, we noticed a formation of ladders consistent with ubiquitination on ^32^P-NAD^+^-Ub (^32^P-NAD^+^-Ub_n_). A previous study reported that RNF114 efficiently catalyses formation of polyubiquitin chains (*34*). Indeed, we also observe that RNF114 forms free polyubiquitin chains in a time- and concentration-dependent manner (**Supplementary** Fig. 5B). We postulate that the ^32^P-NAD^+^-Ub_n_ ladders are generated as RNF114 transfers Ub onto existing ^32^P-NAD^+^-Ub catalysed by DTX2-RD. To verify this hypothesis, we tested both DTX2-RD and RNF114 in the presence of ubiquitin cascade components (E1/E2 (UbcH5B)/Ub/Mg^2+^-ATP) and biotin-Ub or biotin-ADPr-Ub. In reactions with biotin-Ub, we observe auto-ubiquitination with DTX2-RD and formation of Ub_2_, Ub_3_, and Ub_4_ ladders with RNF114 (**Fig. 3B**, first group). In reactions with biotin-ADPr-Ub, we only observe Ub_2_, Ub_3_, and Ub_4_ ladders with RNF114, and no DTX2 autoubiquitination (**Fig. 3B**, second group). RNF114 prefers to catalyse Ub transfer to biotin-ADPr-Ub probe **1**, as opposed to biotin-Ub, as is evident from the remaining unanchored biotin-Ub bands, with 16% remaining ADPr-Ub probe **1** and 70% remaining biotin-Ub after 30 minutes (**Fig. 3C, D**).

**Figure 3.**
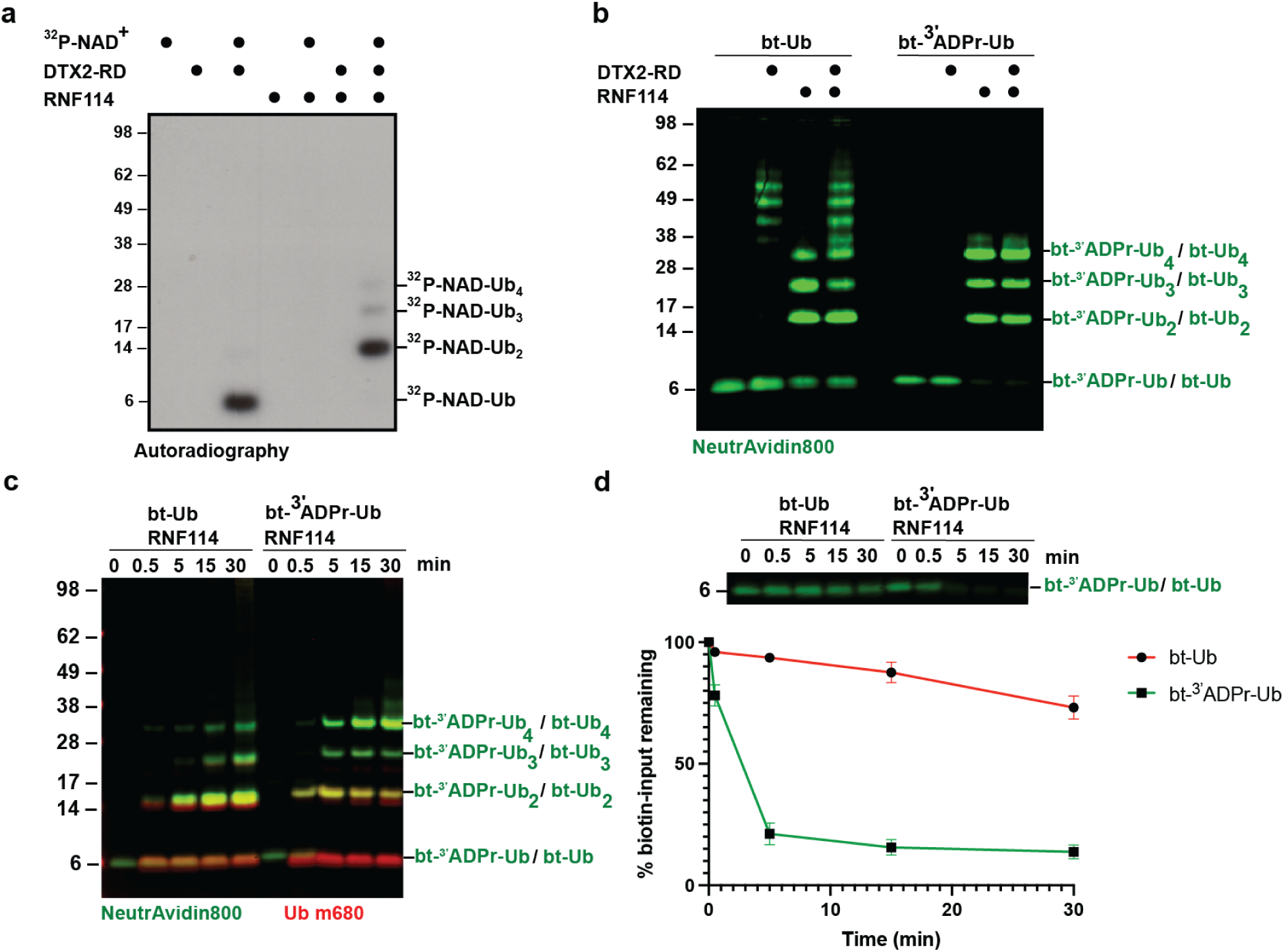
RNF114 elongates ADPr-Ub. **A)** RNF114 elongates Deltex-catalysed ^32^P-NAD^+^-Ub, **B)** RNF114 elongates biotin-Ub and biotin-ADPr-Ub probe **1**, **C)** RNF114 elongates biotin-ADPr-Ub probe in a time-dependent manner. A-C: all in the presence of E1/UbcH5B/Ub/Mg^2+^-ATP. **D)** Quantification of remaining biotin-Ub or biotin-ADPr-Ub probe **1** from C.

### RNF114 elongates ADPr-Ub via K11 linkage

To understand how RNF114 elongates ADPr-Ub, we looked at the Alphafold-predicted structural model of RNF114 and ubiquitin (**Fig. 4A**, left). RNF114 possesses a Ub-Interacting Motif (UIM) domain in its C-terminus, which is predicted to engage with the hydrophobic Ile44 patch of Ub. Interestingly, the Ub-tail in the model is pointing towards the tandem ZnF2+ZnF3 domains of RNF114, which were suggested as mono-ADPr or poly-ADPr reader(*24, 25*). This opens the possibility that RNF114 binds ADPr-Ub through the ZnF2+ZnF3+UIM domains. We then modelled in UbcH5B–Ub (PDB: 4V3L) to bind to the RING domain of RNF114, and in this scenario, the UbcH5B^K85^–Ub^Gly76^ isopeptide bond is poised towards the acceptor Ub^K11^ within a ∼7 Å distance (**Fig. 4A**, right). This model would predict that the polyubiquitin chains formed by RNF114 are of the K11 linkage type. To validate this model, we tested if RNF114 can build specific Ub chain linkage types in an unbiased manner. We utilised a mixture of all eight neutron-encoded Ub acceptors carrying only one amine position (M1, K6, K11, K27, K29, K33, K48, K63)(*35*) and tested which of those acceptors RNF114 can elongate with an all-Lys-to-Arg Ub donor. Out of the eight acceptors, RNF114 consumes Ub K11 most significantly and likewise, mainly K11-linked Ub_2_ (92%) is formed as product over time, with K48-linked Ub_2_ as minor side product (7%) (**Fig. 4B,C, Supplementary** Fig. 6).

**Figure 4.**
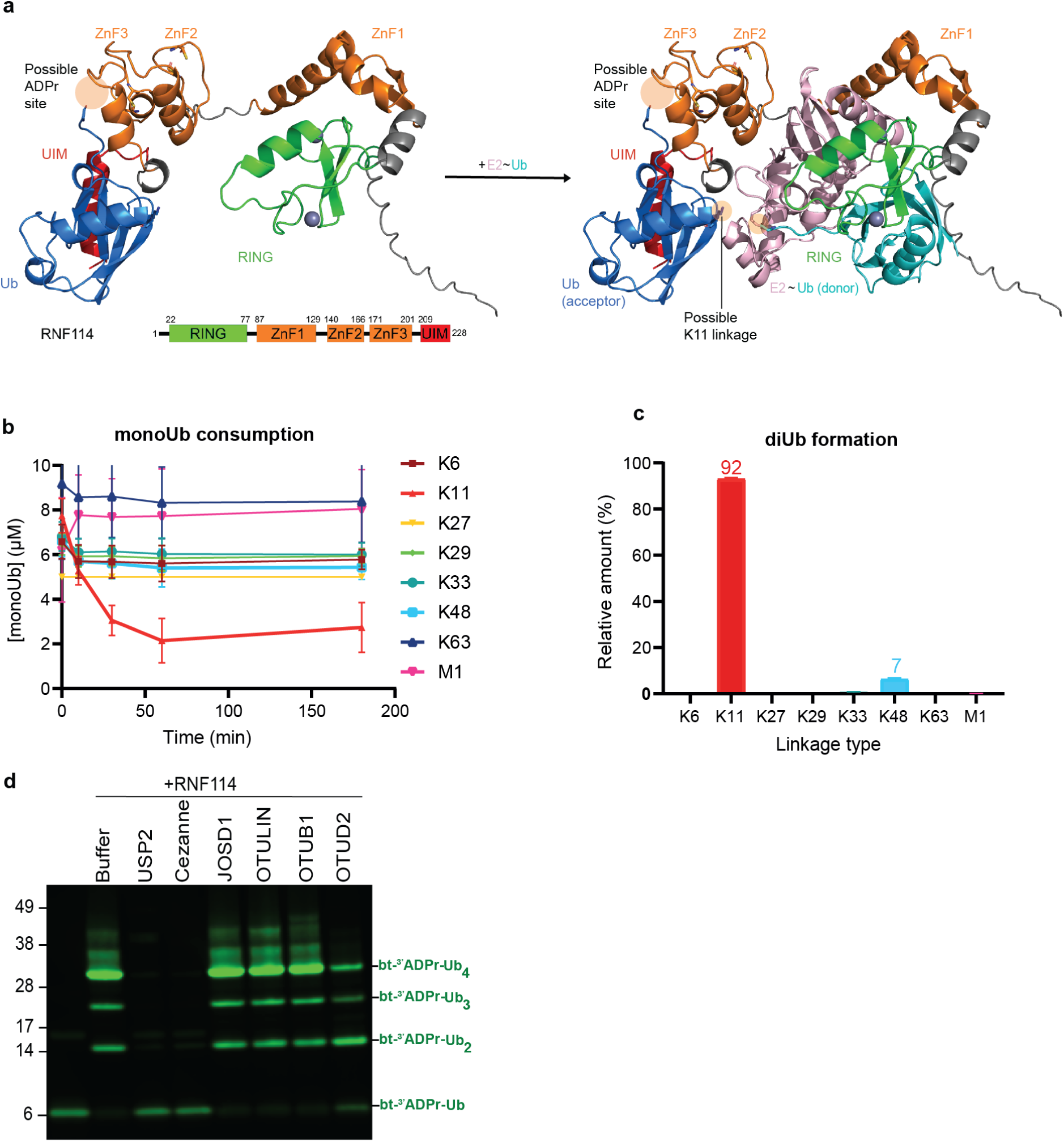
RNF114 elongates ADPr-Ub via K11 linkage. **A)** Structural prediction model of RNF114 bound to acceptor Ub and UbcH5B–Ub. The donor UbcH5B^K85^–Ub^Gly76^ isopeptide bond is poised towards the acceptor Ub^K11^, **B, C)** RNF114 prefers to elongate Ub on K11, as shown by neutron encoded mono-Ub consumption and di-Ub formation, **D)** Panel of DUBs to remove RNF114-catalysed poly-ubiquitinated biotin-ADPr-Ub probe. Cezanne (K11-specific DUB) removes the poly-ubiquitinated species.

We then tested a panel of deubiquitinases (DUBs) for the removal of RNF114-catalysed poly-ubiquitination of biotin-ADPr-Ub probe **1** (**Fig. 4D, Supplementary** Fig. 7). We found that USP2 (a promiscuous DUB) and Cezanne (a K11 linkage specific DUB) were able to remove all biotin-ADPr-Ub*_n_* signal as well as Ub*_n_* chains into mono biotin-ADPr-Ub. and mono Ub. OTUD2 (K11, K27, K29, K33-linkage specific DUB) showed weak removal activities, which reduces biotin-ADPr-Ub*_n_* although not completely. JOSD1, OTULIN (M1 specific DUB), and OTUB1 (a K48 specific DUB) did not show removal in the condition tested. These results are consistent with our unbiased approach using neutron-encoded Ub acceptors and further suggest that RNF114 elongates ADPr-Ub mainly through K11 linkage.

### RNF114 engages ADPr-Ub through its ZnF2+ZnF3+UIM domains

Next, we went on to validate the minimum RNF114 domains required for ADPr-Ub binding. The structural model suggested that RNF114 possibly binds ADPr-Ub through the ZnF2+ZnF3+UIM domains. To test this, we generated RNF114 domain truncations (**Fig. 5A**) and tested their binding to the biotin-ADPr-Ub probe **1** using BLI. The loss of the UIM domain resulted in ∼10x worse affinity, whereas the tandem ZnF2+ZnF3 alone resulted in ∼27x worse affinity compared to the full-length protein. Meanwhile, the ZnF2+ZnF3+UIM domains bind with 1.4 µM affinity, comparable to 850 nM affinity of the full-length protein. Deletion of ZnF2+ZnF3+UIM domains resulted in a complete loss of detectable binding (**Fig. 5B, Supplementary** Fig. 8A). Together, these binding experiments suggest that ZnF2+ZnF3+UIM domains are the minimum module required for decent ADPr-Ub binding by RNF114.

**Figure 5.**
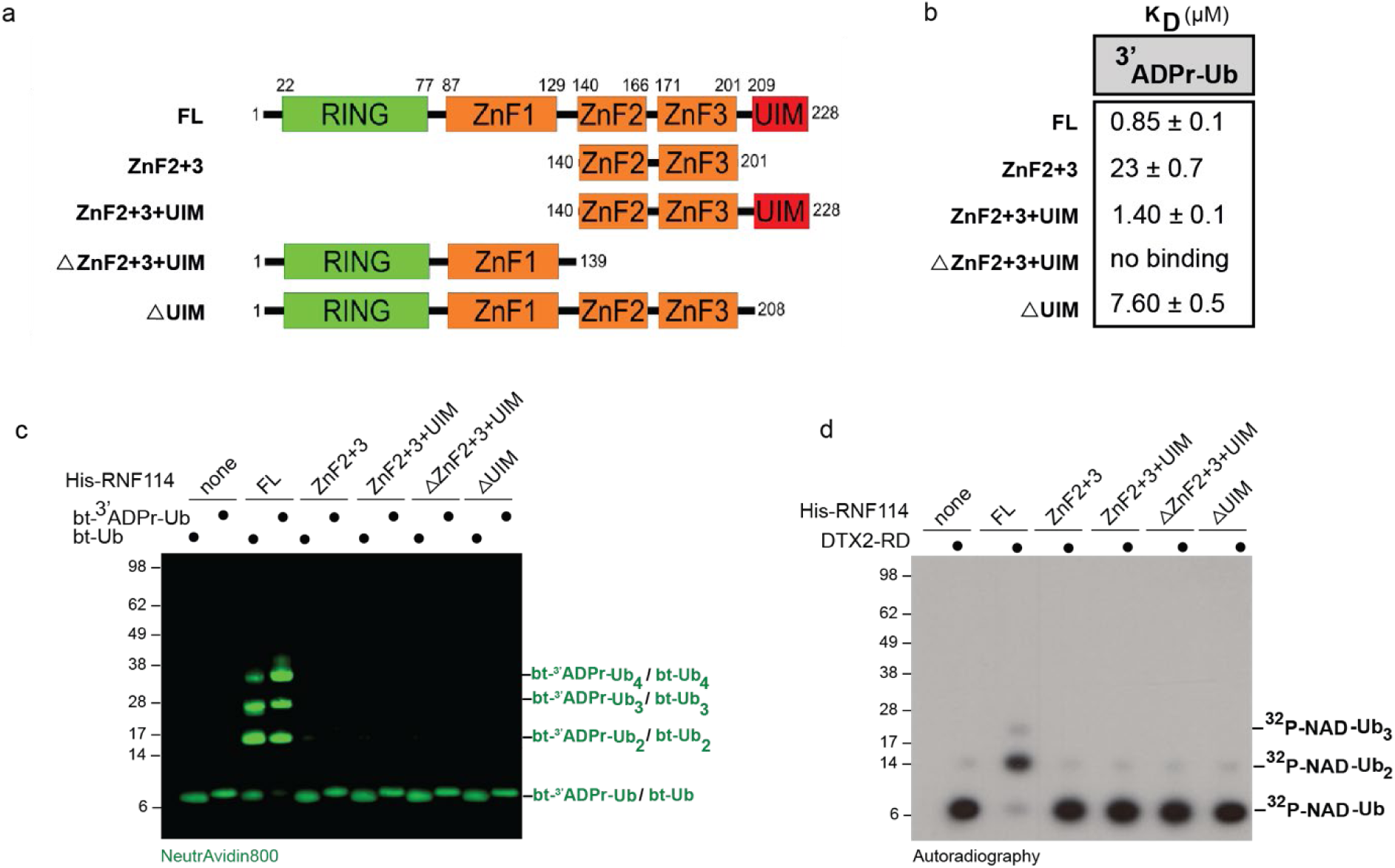
RNF114 ZnF2+ZnF3+UIM domains are responsible for ADPr-Ub binding. **A)** RNF114 domain truncation scheme, **B)** Binding affinities of RNF114 FL and truncation mutants to ADPr-Ub probe **1**, **C)** Biotin-ADPr-Ub probe **1** *in vitro* elongation reactions with RNF114 FL and truncations, **D)** ^32^P-NAD^+^-Ub *in vitro* elongation reactions with RNF114 FL and truncations.

We also tested the RNF114 truncations in Ub and ADPr-Ub elongation reactions. None of the truncations were able to elongate biotin-Ub or biotin-ADPr-Ub (**Fig. 5C, Supplementary** Fig. 8B). In an analogous set of experiments, we tested these truncations using DTX2-dependent ^32^P-NAD^+^-Ub with similar results (**Fig. 5D, Supplementary** Fig. 8C). Even ΔZnF2+3+UIM and ΔUIM variants which still contain the RING domain were unable to elongate Ub or ^32^P-NAD^+^-Ub. This suggests that RNF114 requires both the RING domain and UIM domains to elongate Ub and ADPr-Ub. Taking together these *in vitro* assays and binding experiment results indicate that RNF114 utilises its ZnF2+ZnF3+UIM domains to bind ADPr-Ub, which then elongates the Ub chain using its RING E3 ligase activity.

### RNF114 recruitment to sites of DNA Damage Response sites *in vivo* is dependent on ZnF2+3+UIM domains

Previous studies show that upon DNA damage, RNF114 is recruited to the sites of PARP1-dependent ADP-ribosylation, which can be either mono-ADPr or poly-ADPr-dependent (*24, 25*). Since we identified that ZnF2+3+UIM domains of RNF114 is a ADPr-Ub binding module, we wondered if RNF114 recruitment to DNA damage sites are dependent on these domains. We generated full-length YFP-RNF114 with mutations on the ZnF2+ZnF3 (C176A) and UIM domains (V220G, L221G, S224G) that are predicted to impair ADPr and Ub binding, respectively, and compared their recruitment to laser-induced DNA damage. We found that these mutations significantly impair RNF114 recruitment to the sites of damage (**Fig. 6**). Deleting the UIM domain also abolishes RNF114 recruitment to the damage sites, which suggest that RNF114 localisation to damage sites does not solely depend on ADPr signal, but also relies on Ub signal. These data show that RNF114 relies on both the UIM and ZnF2+ZnF3 domains for its recruitment to DNA damage sites, and that this is likely to be driven by ADPr-Ub signal.

**Figure 6.**
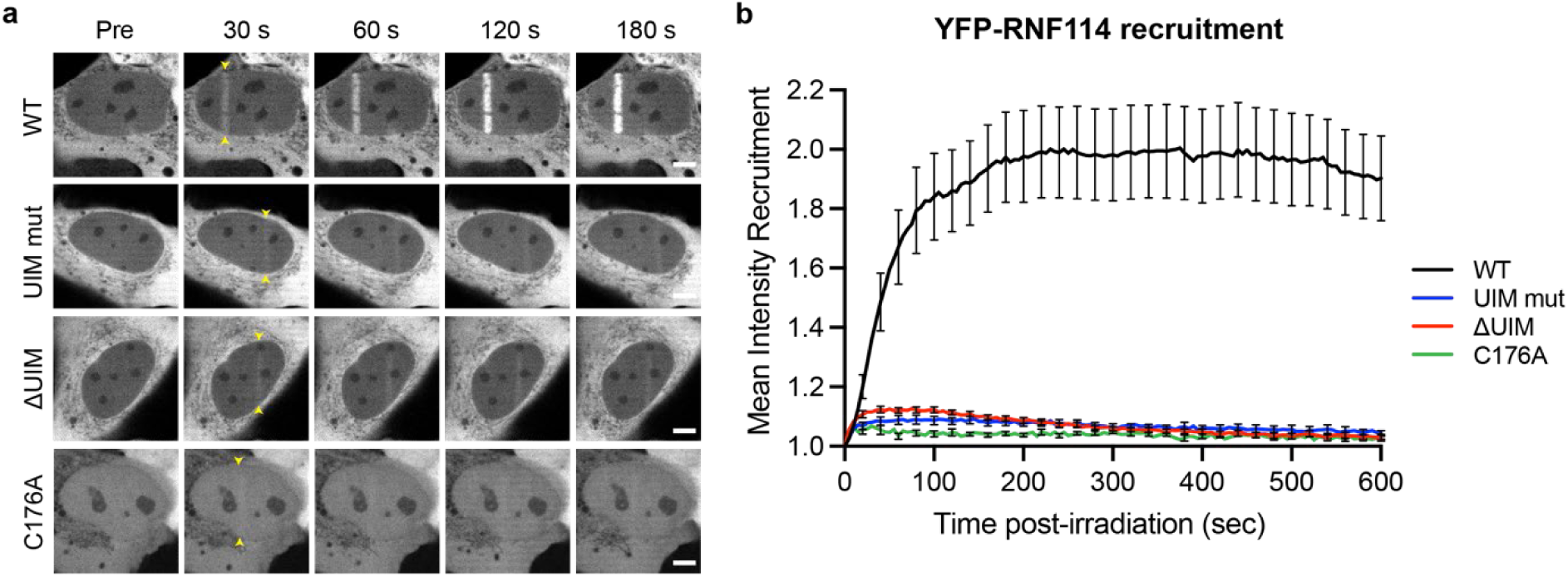
RNF114 recruitment to sites of DNA damage is dependent on its tandem ZnF2+ZnF3+UIM domains. **A)** Representative images of YFP-RNF114 WT or mutants recruitment to sites of DNA damage induced by 405 nm irradiation in U2OS WT cells. Arrow heads indicate site of damage. Scale bar, 5µm, **B)** Recruitment kinetics of YFP-RNF114 WT (Black), UIM mutant (Blue), ΔUIM (Red), or C176A (Green) to sites of damage. Data is representative of two independent replicates. Curves show mean ± s.e.m. of at least 14 cells per condition.

## Discussion

Ubiquitination of non-protein substrates is an emerging theme. However, investigating their biological function is challenging due to the lack of tools. The synthesis of non-hydrolysable ADPr-Ub analogue **1** allowed us to identify RNF114 in an unbiased manner as a reader of this recently discovered modification. The tandem ZnF2+ZnF3+UIM domains of RNF114 bind the ADPr-Ub probe with an affinity below 1 µM. This is surprisingly high, considering that UIM domains typically bind Ub with low affinity, with K_D_ > 100 µM(*36*). For Ub-binding domains, tighter binding and linkage selectivity is usually achieved through the arrangement of multiple domains to achieve linkage-specific avidity(*37*). Our data suggests that RNF114 achieves tight ADPr-Ub binding due to having adjacent ADPr-binding ZnF2+ZnF3 and Ub-binding UIM domains. It will be interesting to see if there are other proteins with adjacent ADPr-reader and Ub-reader functions that confer similar ADPr-Ub binding ability.

RNF114 is an E3 ligase, and it contains a C3HC4-type RING finger domain to support this function. Our biochemical reconstruction and structural modelling demonstrate that the ADPr-Ub binding modules cooperate with the RING domain to produce a novel mixed Ub linkage product, K11-linked ADPr-Ub*_n_*. Also, the ability of RNF114 to elongate ADPr-Ub fits nicely with recent findings observed by other research groups. For example, DTX1/2/4 were reported to interact with PARylated TNKS1, which led to the stabilisation of mono-Ub TNKS1. RNF114 and its paralogue RNF166 interact with this mono-Ub TNKS1 and promotes its K11-linked di-ubiquitination(*32*). In particular, the tandem ZnF2+ZnF3+UIM domains were sufficient to stabilise mono-Ub TNKS1 species; these are the same domains identified in our study as ADPr-Ub binder. Our findings offer an explanation to complement these observations: that the DTX1/2/4-dependent mono-Ub TNKS1 species is possibly TNKS1-(^3’^ADPr-Ub). Another recent study reported ubiquitination on mono-ADP-ribosylated PARP10, and shows that this can be extended with K11-linked chains(*38*). It is highly possible that this K11-linked elongation is mediated by RNF114 or other paralogues found in human cells such as RNF166. These two examples indicate that ADPr-Ub modification on a substrate are uniquely recognised and have distinct biological outcomes. It remains to be seen if there are other possible substrates and fates for ADPr-Ub modification besides K11-linked elongation by RNF114.

Previously, it was shown that RNF114 is recruited to DNA damage in an ADP-ribosylation dependent manner and could be either mono-ADPr or poly-ADPr-dependent(*24, 25*). Another recent report showed that RNF114 recruitment to the sites of DNA damage is dependent on its UIM domain (*39*). Excitingly, our data demonstrate that RNF114 recruitment to DNA damage sites rely on both the ZnF2+ZnF3 and UIM domains. Paired with our identification of RNF114 as a tight ADPr-Ub binder, this strongly suggests that the UIM and ZnF2+ZnF3 domains act as a specific dual reader module for the ADPr-Ub signal. It is known that PAR and MAR signals are heavily induced upon DNA damage, which are ubiquitination targets of Deltex E3 ligases (*10, 12, 13*). Moreover, ADP-ribosylation-dependent ubiquitination was also observed under interferon stimulation and the signal is dependent on both PARP14 and DTX3L/PARP9 (*18*), the latter complex being an ADPr-Ub writer. This suggests that there might be a higher level of ADPr-Ub-substrate modification under such conditions. Although more labile compared to lysine ubiquitination, ester-linked ubiquitination has a cellular half-life of at least 1 hour (*40*) in which the signal has the opportunity to be recognised and potentially drive biological outcomes.

In this work, we synthesised a non-hydrolysable ADPr-Ub analogue and identified RNF114 as the first reader of the dual ADPr-Ub modification. The development of chemical tools, such as Ub-PAR analogues, might facilitate the identification and characterisation of other readers of these non-canonical Ub codes. Our findings also open up the possibility to utilise RNF114 ZnF2+ZnF3+UIM domains to stabilise endogenous ADPr-Ub-substrates and identify them. For example, by using this as an enrichment strategy to preserve the labile ester-linked ADPr-Ub modification and combine it with an ester-compatible proteomics workflow (*26*) to identify ADPr-Ub-substrates and the sites of modification by MS. We also showed that the ADPr-Ub signal can be elongated by an enzyme that possesses both the reader module and the catalytic RING domain, like RNF114. This reveals the possibility of further modification on existing ADPr-Ub signal on a substrate. The recruitment of RNF114 to DNA damage sites suggests that ADPr-Ub may be a novel signalling molecule in DNA damage response. On a related note, interferon stimulation induces ubiquitination foci that are dependent on ADP-ribosylation (*18, 41*). Studying the ADPr-Ub signal under these biological conditions will potentially uncover novel pathways in eukaryotic cells.

## Methods

### Cell Culture

HEK293T (Cat# ATCC® CRL-3216™) cells were cultured in Dulbecco’s modified Eagle’s medium (DMEM) (Gibco). Medium was supplemented with 8% FCS. Cells were grown at 37 °C in a humidified 5% CO_2_ incubator. All cell lines have been routinely tested for mycoplasma.

### Cloning and plasmids

The constructs described in this paper were generated using standard polymerase chain reaction and verified by Sanger sequencing. For *E. coli* expression, RNF114 were cloned into pRSF Duet vector with an N-terminal 6x-His tag followed by a TEV cleavage site. Mutants were generated using QuikChange mutagenesis. For laser irradiation experiment, RNF114 variants with flanking attB1/attB2 sequence were cloned into gateway cloning-compatible system (pDONR221 and pDEST N-terminal YFP).

### Protein expression and purification

Proteins were expressed in *E. coli* Rosetta (DE3) cells. Cultures were grown in LB medium with appropriate antibiotic at 37 °C until they reach an OD_600_ of 0.6-0.8, then protein expression was induced by supplementing 0.2 mM isopropyl-β-D-1-thiogalactopyranoside (IPTG) at 18 °C for 16-18 hours. Cells were harvested and resuspended in buffer A (50 mM Tris-HCl pH 8.0, 200 mM NaCl, 20 mM imidazole, 5 mM β-mercaptoethanol) and supplemented with 0.2 mM phenylmethylsulfonyl fluoride (PMSF). The resuspended cells were lysed using homogeniser at 15,000 psi and cleared by high-speed centrifugation at 17,000 rpm for 1h at 4 °C. The cleared lysate was incubated with Ni^2+^-NTA resin or HisTrap FF column (Cytiva), washed in buffer A, and eluted in buffer B (50 mM Tris-HCl pH 7.5, 200 mM NaCl, 200 mM imidazole, 5 mM β-mercaptoethanol). Eluted proteins were further purified by size exclusion chromatography on a Superdex 200 column (Cytiva) pre-equilibrated in 20 mM HEPES pH 7.5, 150 mM NaCl, 1 mM DTT. Fractions of interest were checked on SDS-PAGE, pooled, and concentrated on Amicon protein concentrator device (Merck Millipore). Protein concentrations were determined by A280 measurement using a nanodrop device and calculated using predicted molar extinction coefficients. Proteins were kept in aliquots, snap-frozen in liquid nitrogen, and stored in −70 °C. His-E1, UbcH5B, His-DTX2-RD, USP2, Cezanne, JOSD1, OTULIN, OTUD2, OTUB1 were generated as previously described.

### In vitro assays

#### RNF114 Ub chain formation assay

Reactions were performed in 50 mM Tris-HCl pH 8.0, 50 mM NaCl, 2.5 mM MgCl_2_, 2.5 mM ATP, 0.2 µM His-E1, 2 µM UbcH5B, 20 µM Ub, and the indicated RNF114 concentration at 37 °C for up to 1 hour. Reactions were stopped by adding 4x LDS and 100 mM DTT. Samples were loaded onto SDS-PAGE (4-12% Bis-Tris gel/MES buffer) and visualised by Instant Blue stain.

#### 32P-NAD+-Ub elongation assay

Reactions were performed in 50 mM Tris-HCl pH 8.0, 50 mM NaCl, 2.5 mM MgCl_2_, 2.5 mM ATP, 0.2 µM His-E1, 2 µM UbcH5B, 20 µM Ub, 1 µM DTX2-RD, and 40 µM NAD^+^ spiked with 40 nCi/µL ^32^P-NAD^+^ at 37 °C for 30 minutes to generate ^32^P-NAD^+^-Ub. To initiate elongation, 1 µM His-RNF114 (FL or variants) was added and incubated at 37 °C for 30 minutes. Reactions were stopped by adding 4x LDS and 100 mM DTT. Samples were loaded onto SDS-PAGE (4-12% Bis-Tris gel/MES buffer) and visualised by autoradiography.

#### Biotin-ADPr-Ub analogue elongation assay

Reactions were performed in 50 mM Tris-HCl pH 8.0, 50 mM NaCl, 2.5 mM MgCl_2_, 2.5 mM ATP, 0.2 µM His-E1, 2 µM UbcH5B, 20 µM Ub, 10 µM biotin-ADPr-Ub, and 1 µM His-RNF114 (FL or variants) at 37 °C for 30 minutes. For Fig 3B-C, either 10 µM biotin-Ub or biotin-ADPr-Ub were added as indicated. For samples treated with DUBs (Fig 4D), reactions were stopped with 50 mM EDTA, followed by addition of 1 µM DUBs and further incubation at 37 °C for 30 minutes. Reactions were stopped by adding 4x LDS and 100 mM DTT. Samples were loaded onto SDS-PAGE (4-12% Bis-Tris gel/MES buffer), transferred onto nitrocellulose membrane (Biorad TransBlot Turbo), and blocked with 5% BSA. The membrane was incubated with primary antibodies (mouse monoclonal anti-Ub(PD41), Santa Cruz Biotechnology sc-8017) overnight at 4°C followed by secondary Li-Cor goat-anti mouse IRDye 680 and NeutrAvidin800 antibodies (Thermo Fisher). Membranes were washed in 0.1% PBS-Tween 20 (PBST) and scanned using Li-Cor Odyssey CLx imager.

#### Modelling

Structural prediction of RNF114 bound to Ub and E2 UbcH5B were obtained using AlphaFold2 Multimer/ColabFold and AlphaFold3 (*42–44*). The donor Ub was modelled in using a structure of UbcH5B–Ub bound to RNF38 (PDB: 4V3L), by aligning the Cα of both UbcH5B (RMSD= 0.68 over 144 residues). The resulting model shows the UbcH5B^K85^–Ub^Gly76^ isopeptide bond is poised towards the acceptor Ub^K11^ within a ∼7 Å distance. Structural figures were prepared in PyMOL (Schrodinger).

#### SDS-PAGE and Immunoblotting

Samples were resolved on precast Bis-Tris NuPAGE gels (Invitrogen, 4-12%, 12%). MOPS buffer (Invitrogen Life Technologies, Carlsbad, USA) for Bis-Tris gels was used as running buffer. Proteins were transferred to a nitrocellulose membrane (Protan BA85, 0.45 μm, GE Healthcare) at 300 mA for 2.5 h. The membranes were blocked in 5% milk (skim milk powder, LP0031, Oxiod) in 1× PBS (P1379, Sigma-Aldrich), incubated with a primary antibody diluted in 5% milk in 0.1% PBS-Tween 20 (PBST) overnight, washed three times for 10 min in 0.1% PBST, incubated with the secondary antibody diluted in 5% milk in 0.1% PBST for 30-45 min, and washed three times again in 0.1% PBST. Immunoblot signals were visualized with a LI-COR Odyssey Fx laser scanning fluorescence imager and analyzed using LI-COR Image Studio Lite software.

#### Antibodies

Primary antibodies used for the study were: mouse anti-DTX2 (ProteinTech, #67209-1-Ig), rabbit anti-RNF114/ZNF313 (Invitrogen, #PA5-98120), mouse anti-Ub (Santa Cruz Biotechnology, sc-8017). Secondary antibodies used were: IRDye 800CW goat anti-rabbit IgG (H + L) (Li-COR, Cat# 926-32211, 1:5000), IRDye 800CW goat anti-mouse IgG (H + L) (Li-COR, Cat# 926-32210, 1:5000).

#### Pull-down assays

For pull-down, the medium was completely removed from the cells, followed by the addition of 300 µL lysis/co-IP buffer (50 mM TRIS, 150 mM NaCl, 0.8% NP40, Roche Protease inhibitor) and scraping of the cells. Cell lysis was done as described above. The supernatant was transferred to a new Eppendorf tube. 15-20 µL of supernatant was removed to blot for the input sample. The remaining supernatant was incubated with a probe (diluted in the same buffer to a final concentration of 5 µM) for 60 mins at 37 °C. Next, the total volume was completed to 1 mL with the lysis buffer. 30 µL of Neutravidin beads (Thermo Scientific™, Pierce™ NeutrAvidin™ Agarose, Catalog number: 29200) were added, followed by incubation at 4 °C on rollers overnight. Beads were collected by centrifugation at 500xg for 1 min at 4 °C and washed with wash buffer (50 mM TRIS, pH 7.5, 150 mM NaCl, 0.5% Triton X, 0.5% SDS) at least 3 times by centrifuging at 500xg for 1 min at 4 °C. Prior to the final washing, beads were transferred to a new tube to avoid background and finally centrifuged at 500xg for 3 mins at 4 °C. Beads and supernatant were stored at −80 °C until following analyses, including immunoblotting, fluorescent scanning, Coomassie staining and LC/MS-MS.

#### Processing of protein gel bands

For MS analysis, gel slices were washed, subjected to reduction with dithiothreitol, alkylation with iodoacetamide and in-gel trypsin digestion using a Proteineer DP digestion robot (Bruker). Tryptic peptides were extracted from the gel slices and lyophilized.

#### Mass spectrometry

Peptides were dissolved in water/formic acid (100/0.1 v/v) and analyzed by on-line C18 nanoHPLC MS/MS with a system consisting of an Ultimate3000nano gradient HPLC system (Thermo, Bremen, Germany), and an Exploris480 mass spectrometer (Thermo). Samples were injected onto a cartridge precolumn (300 μm × 5 mm, C18 PepMap, 5 um, 100 A, and eluted via a homemade analytical nano-HPLC column (50 cm × 75 μm; Reprosil-Pur C18-AQ 1.9 um, 120 A (Dr. Maisch, Ammerbuch, Germany). The gradient was run from 2% to 40% solvent B (20/80/0.1 water/acetonitrile/formic acid (FA) v/v) in 30 min at 250 nl/min. The nano-HPLC column was drawn to a tip of ∼10 μm and acted as the electrospray needle of the MS source. The mass spectrometer was operated in data-dependent MS/MS mode, with a HCD collision energy at 30% and recording of the MS2 spectrum in the orbitrap, with a quadrupole isolation width of 1.2 Da. In the master scan (MS1) the resolution was 120,000, the scan range 400-1500, at standard AGC target and a maximum fill time of 50 ms. A lock mass correction on the background ion m/z=445.12003 was used. Precursors were dynamically excluded after n=1 with an exclusion duration of 10 s, and with a precursor range of 20 ppm. Included charge states were 2-5. For MS2 the first mass was set to 110 Da, and the MS2 scan resolution was 30,000 at an AGC target of 100% at a maximum fill time of 60 ms. Maxquant version 2.5.1.0 was used for label free data processing with 10 ppm and 0.02 Da deviation for precursor and fragment mass, respectively. Trypsin was specified as the enzyme. Methionine oxidation and the acetylation (on the protein N-terminus) were set as variable modifications. Carbamidomethyl was set as a fixed modification on cysteines. The false discovery rate was set < 1%.

#### Bioinformatic analysis of DIA data

Peptide intensity table with the Label-free quantitation (LFQ) values was analyzed in Perseus (v1.6.2.3). Data were log_2_ transformed and filtered for identification in all three replicates in at least one group. Principal component analysis (PCA) was performed for each analysis with default settings. Intensities were *Z-scored* by subtracting the mean and used for hierarchical clustering by Euclidean distance (pre-processed with k-means, 300 clusters, 1000 iterations). Missing values were imputed from the lower end of the normal distribution (default settings). A two-sided student’s t-test with permutation-based FDR was used to calculate the significance between treatment and control at 0.05 FDR (p-value). For a detailed protocol of data analysis in Perseus: https://cox-labs.github.io/coxdocs/interactions.html.

#### Data availability

The mass spectrometry proteomics data has been deposited to the ProteomeXchange Consortium via the PRIDE partner repository with the dataset identifier *PXD58087*.

### Bio Layer Interferometry (BLI)

#### DTX2 assay

Bio Layer Interferometry (BLI) measurements were conducted using an OctetRed system (ForteBio). Streptavidin-biosensor were equilibrated in buffer (25 mM Tris-Cl 100 mM NaCl, BSA (1 mg/mL), dextran (1 mg/mL), Tween 20 (0.05%) and DTT (1 mM) pH 7.5,) before loaded with Bt-ADPr-Ub (**1**), or the controls Bt-Ub (no ADPr), or Bt-ADPr (no Ub). The sensor tips were then transferred to solutions of DTX2 (ranging from 40 µM - 0.163 µM) to assess the association rates of the analyte. Subsequently, the sensor tips were returned into buffer to reverse the ligand-protein interaction and estimate the corresponding dissociation rates. Dissociation constants (K_D_) were then computed using the ForteBio Data Analysis software, employing simultaneous co-fitting of all concentrations.

#### RNF114 or RNF114 domain deletions assays

Bio Layer Interferometry (BLI) measurements were conducted using an OctetRed system (ForteBio). Streptavidin-biosensor were equilibrated in buffer (50 mM HEPES, 50 mM NaCl, 5 mM MgCl_2_, 1 mM DTT, BSA (1 mg/mL), dextran (1 mg/mL) and Tween 20 (0.05%) ,pH 7.5) before loaded with Bt-ADPr-Ub (**1**) or the controls Bt-Ub (no ADPr), or Bt-ADPr (no Ub). The sensor tips were then transferred to solutions of RNF114 or RNF114 domain deletions (ranging from 22 µM - 0.83 µM) to assess the association rates of the analyte. Subsequently, the sensor tips were returned into buffer to reverse the ligand-protein interaction and estimate the corresponding dissociation rates. Dissociation constants (K_d_) were then computed using the ForteBio Data Analysis software, employing simultaneous co-fitting of all concentrations.

### *In vitro* conjugation assays with mass spectrometry read-out

To test the chain-building capacity and chain-building specificity of RNF114, an assay with neutron-encoded monoubiquitins was used.^35^ In short, eight neutron-encoded monoubiquitin_1-75_ acceptors (with only one available lysine position, all other lysine were mutated —to-R) were mixed in an equimolar amount (8x ∼7.5 Μm; 1.5x final concentration) together with donor mono ubiquitin_1-76_ (all K-to-R) (60 µM; 1.5x final concentration). Recombinant purified conjugation enzymes were diluted to 10x final concentration (1 μM for UBE1, 25 μM for UBE2D1 enzyme and 50 µM for RNF114) in a buffer containing 20 mM Tris·HCl, 100 mM NaCl, pH 7.55 and 50 mM TCEP and MgCl_2_. The monoUb mixture (6.33 μL) was added to an Eppendorf tube. Subsequently, the enzymes (1 μL E1, 1 μL E2 and 1 μL E3, 10x final concentration) were added to the monoUb mixture. The mixture was diluted with 0.7 µL buffer (20 mM Tris·HCl, 100 mM NaCl, pH 7.55 and 50 mM TCEP and MgCl_2_). The reaction was started by the addition of 0.2 µL 0.5 M ATP. The reaction mixtures were incubated at 37°C for 180 min. After 30 and 60 minutes fresh ATP (0.2 µL 0.5 M) was added to the solution. Before adding ATP, 1 µL of reaction mixture was taken for timepoint 0 minutes. After 10, 30, 60, and 180 min a sample was taken from the reaction mixture for analysis. After the indicated incubation time, 1 μL of the reaction mixture was taken. The reaction was quenched by acidification of the mixture and internal standards for MS analysis were directly added. Samples were collected in a 96-well plate and measured by LC-MS analysis. 8 μL of the sample was injected onto the column. The samples were separated by an Acquity H-class UPLC system using a BEH C4 column (300Å, 1.7 μM (2.1 x 100 mm)). Proteins were eluted using a shallow gradient that ranged from 29% up to 32% ACN in H_2_O with 0.1% FA over 2 min using a flow rate of 0.6 mL/min, which was able to separate monoUb and diUb products (baseline level). Products were analysed by intact MS analysis on a XEVO G2 XS Q-TOF. The data was analysed using LaCyTools version 22.04.29. Data was aligned, quality control was performed (Mass Accuracy < 15 ppm; IPQ < 0.25; S/N > 9) and the areas of all m/z peaks, within the quality control boundaries, from the same analyte were summed. The absolute area for each monoUb analyte was normalised at each timepoint using the non-processed K27 acceptor monoUb. The concentration of monoUb present at each timepoint was calculated using the theoretical concentration of present K27 acceptor monoUb in the analysed mixture. For the diUb signals, the total absolute areas under the curve for timepoint t= 180 min were summed and the contribution per diUb to this total area was calculated as an indication for the relative amount of that specific diUb linkage type formed during the assay. Results were plotted using GraphPadPrism 10.2.3. ((Code availability. LaCyTools is freely available for download at https://github.com/Tarskin/LaCyTools.))

### Imaging

Cell culture and live-cell imaging was completed as previously described (*45*). Briefly, human U2OS osteosarcoma (American Type Culture Collection (ATCC), HTB-96) cells were cultured in Dulbecco’s modified Eagle’s medium (Sigma-Aldrich) supplemented with 10% fetal bovine serum (FBS; Gibco) and penicillin-streptomycin (100 U/ml; Gibco) in a humidified atmosphere at 37°C with 5% CO2. Cells were plated on an eight-well μ-slide glass bottom chamber slide (ibidi) and transfected with an expression plasmid for YFP-RNF114 WT, YFP-RNF114 UIM mutant (V220G,L221G,S224G), YFP-RNF114 ΔUIM (1-208), or YFP-RNF114 ZnF2+ZnF3 mutant C176A together with PaTagRFP-H2B using TransIT-LT1 Transfection Reagent (Mirus Bio), according to the manufacturer’s instructions. Cells were incubated for 48 hours prior to imaging. For cell sensitization before laser irradiation at 405 nm, growth medium was aspirated from the chamber slide and replaced with fresh medium containing Hoechst 33342 (0.3 μg/mL) for 1 h. Immediately before imaging, the Hoechst containing medium was replaced with imaging media (phenol red–free Leibovitz’s L-15 medium (Life Technologies) supplemented with 20% FBS, penicillin (100 μg/mL), and streptomycin (100 U/mL)). Live-cell microscopy was carried out on an Olympus IX-83 inverted microscope equipped with a Yokogawa SoRa superresolution spinning-disk head, a UPlanAop 60x/1.5 numerical aperture oil-immersion objective lens and a Prime BSI scientific complementary metal-oxide semiconductor camera. The fluorescence of YFP and PaTagRFP-H2B was excited with 488-nm and 561-nm solid state lasers, respectively, and fluorescence detection was achieved with band-pass filters adapted to the fluorophore emission spectra. Laser microirradiation at 405 nm was performed along a 15-μm line through the nucleus for 250 ms using a single-point scanning head (Olympus cellFRAP) coupled to the epifluorescence backboard of the microscope. To ensure reproducibility, laser power at 405 nm was measured at the beginning of each experiment and set to 110 μW at the sample level. For time course experiments, images were collected every 5 s. For the live-cell imaging experiments, cells were maintained at 37°C with a heating chamber. Protein accumulation at sites of damage (*Ad*) was then calculated as:

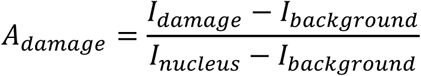

The accumulated protein at the damage (Ad) is a measure of the intensity (I) at the sites of damage over the intensity of the entire nucleus, with subtracted background intensity. The intensity within the microirradiated area was then normalized to the intensity before damage induction. Photoactivated H2B was used as a reference to indicate where irradiation had occurred.

### Statistical analysis

All statistical evaluations were reported on Student’s *t* test (two-tailed distribution) having ns: not significant. All error bars correspond to the mean ± SD. Data were analyzed using the Microsoft Excel and GraphPad Prism5 software.

## Acknowledgments

We would like to thank Alan Wainman and the Dunn School Bioimaging Facility for expert advice and access to the microscope.

