## Supplementary material for "Identification of RNF114 as ADPr-Ub reader through non-hydrolysable ubiquitinated ADP-ribose": Sup. Info

#### Supplementary figures

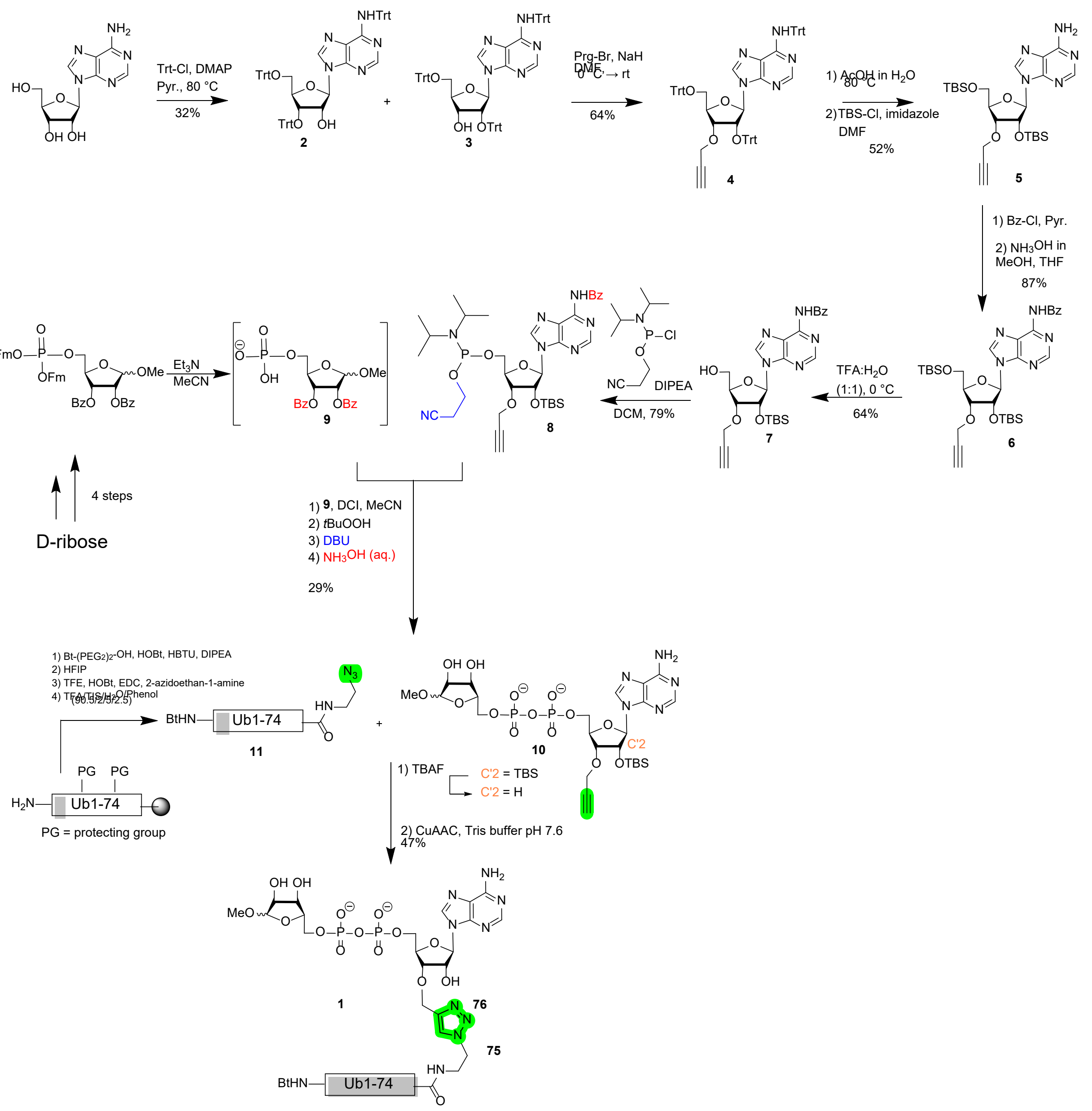

**Supplementary Fig. 1. Synthesis of probe 1.**

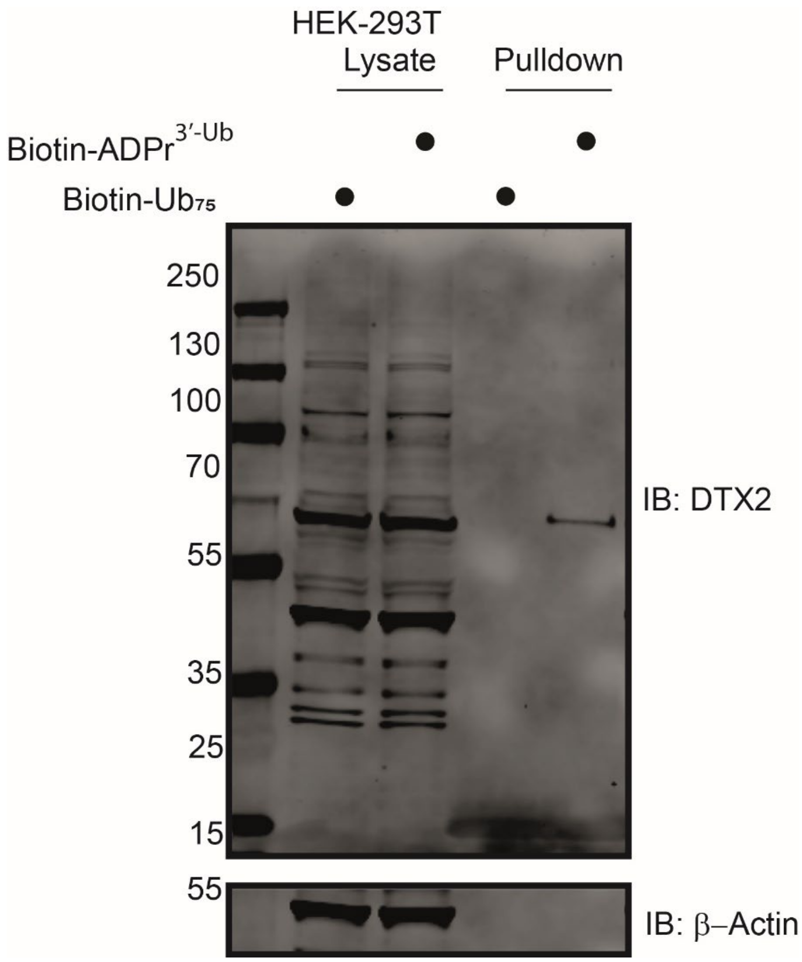


**Supplementary Fig. 2.**

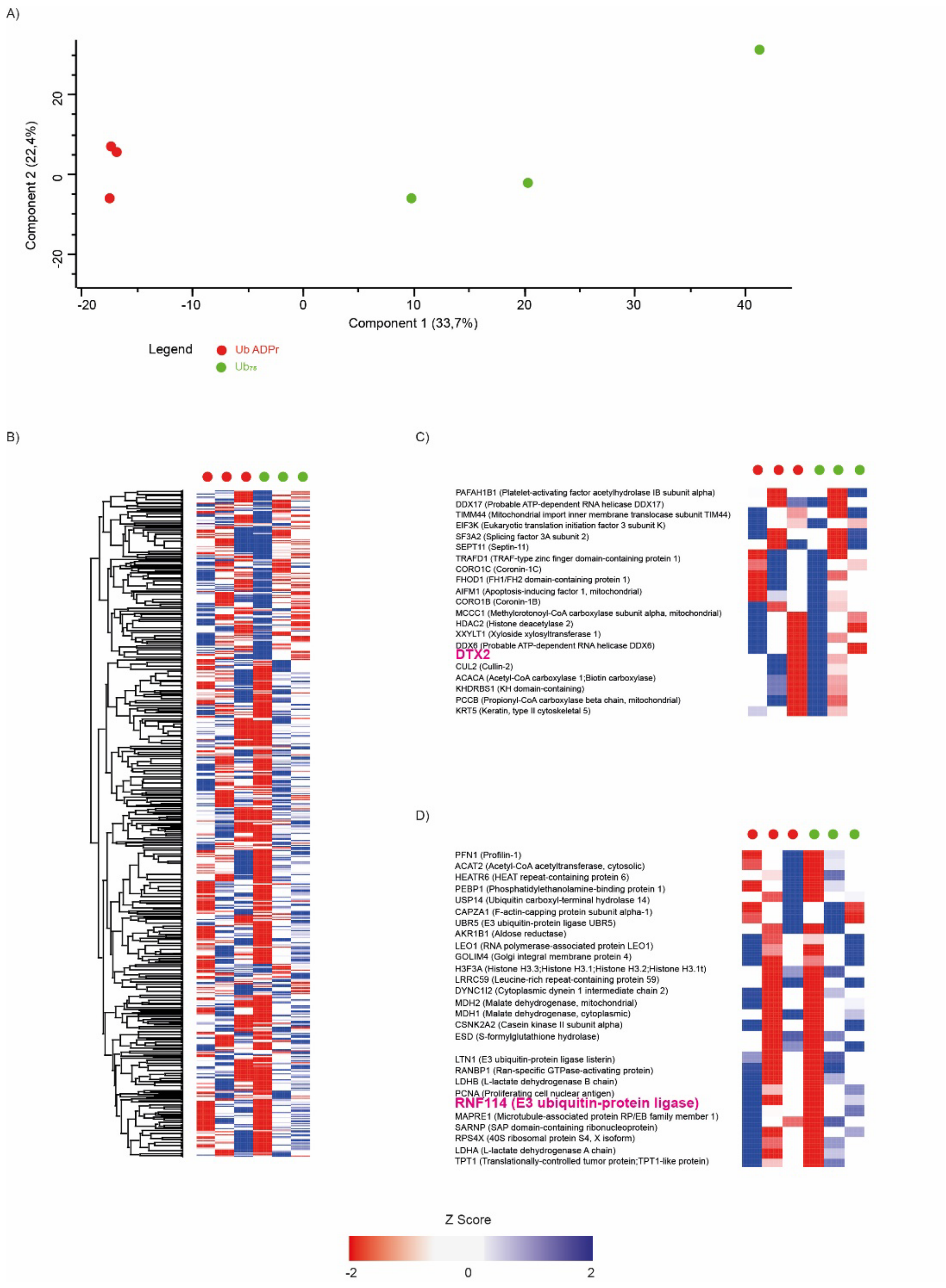


**Supplementary Fig. 3. Principal Component Analysis (PCA) profile and Hierarchical clustering of from probe interactome.** (A) PCA profile of the probe/negative control interactome from the data-independent acquisition (DIA) data using components with the highest explained variance, n = 3. Colored dots indicated represent cell conditions. (B) Heat map showing the hierarchical clustering of the significantly affected genes (p-value < 0.05) from the Total proteome data set. Clustering was done based on Euclidean distance (pre-processed with k-means, 300 clusters, 1000 iterations), where each column in the heatmap is a sample and each row is a protein. The LFQ intensity values were normalized by Z score. Heat map showing the hierarchical clustering of (C) DTX2 and (D) RNF114 highlighted in pink.

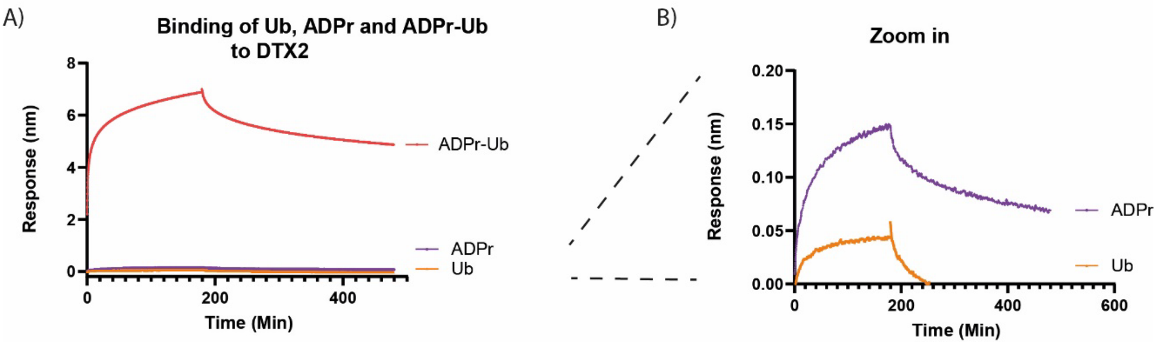


**Supplementary Fig. 4**. BLI measurement comparing the $K_{on}$ and $K_{off}$ rates of $^{3'}$ADPr-Ub **1** to the Bt-Ub and Bt-ADPr (**29**) controls. The graph on the right is a zoomed in version of the left panel, showing minor association rates for Ub and ADPr **29** at 40 μM of DTX2. Therefore, $K_d$ Ub >>> 40 μM and $K_d$ ADPr **29** >> 40 μM.

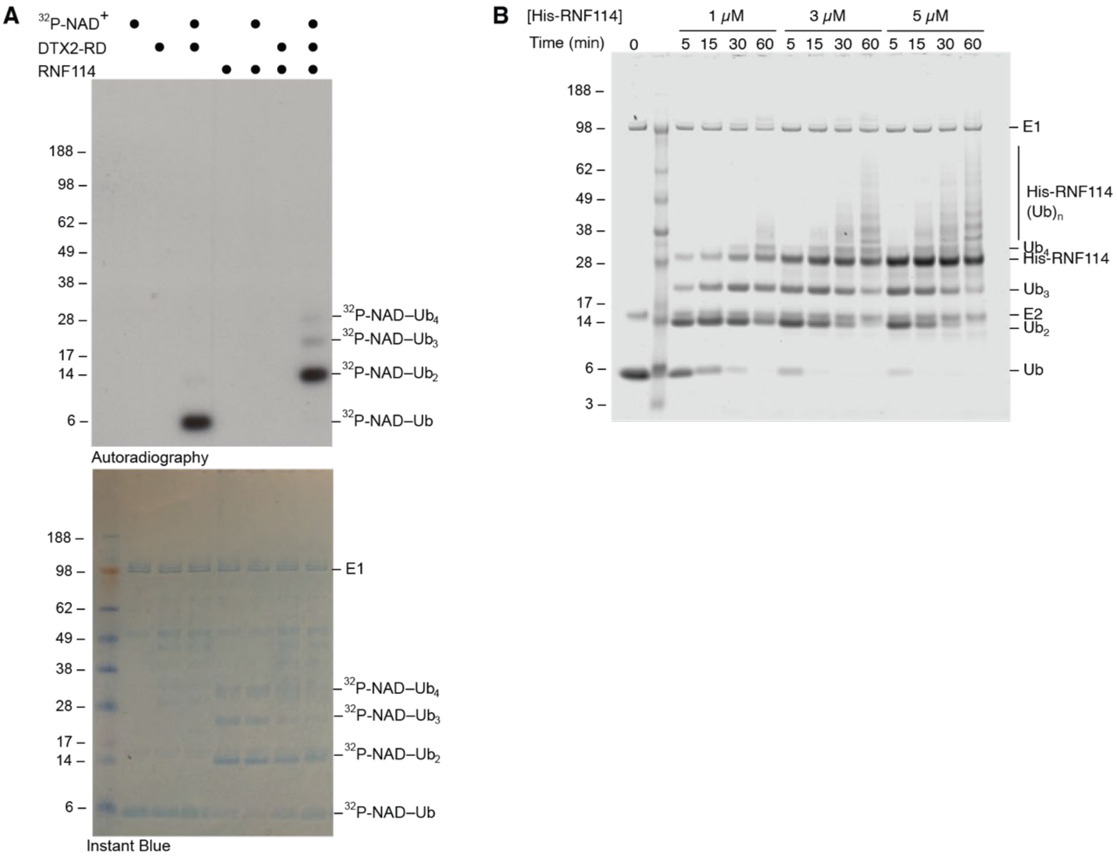


**Supplementary Fig 5. RNF114 elongates $^{3'}$ADPr-Ub. A)** RNF114 elongates $^{32}$P-NAD$^+$–Ub in the presence of E1, E2 (UbcH5B), Ub, $Mg^{2+}$, ATP and DTX2-RD, **B)** RNF114 catalyses free Ub chain formation in the presence of E1, E2 (UbcH5B), Ub, $Mg^{2+}$, ATP in a time dependent manner.

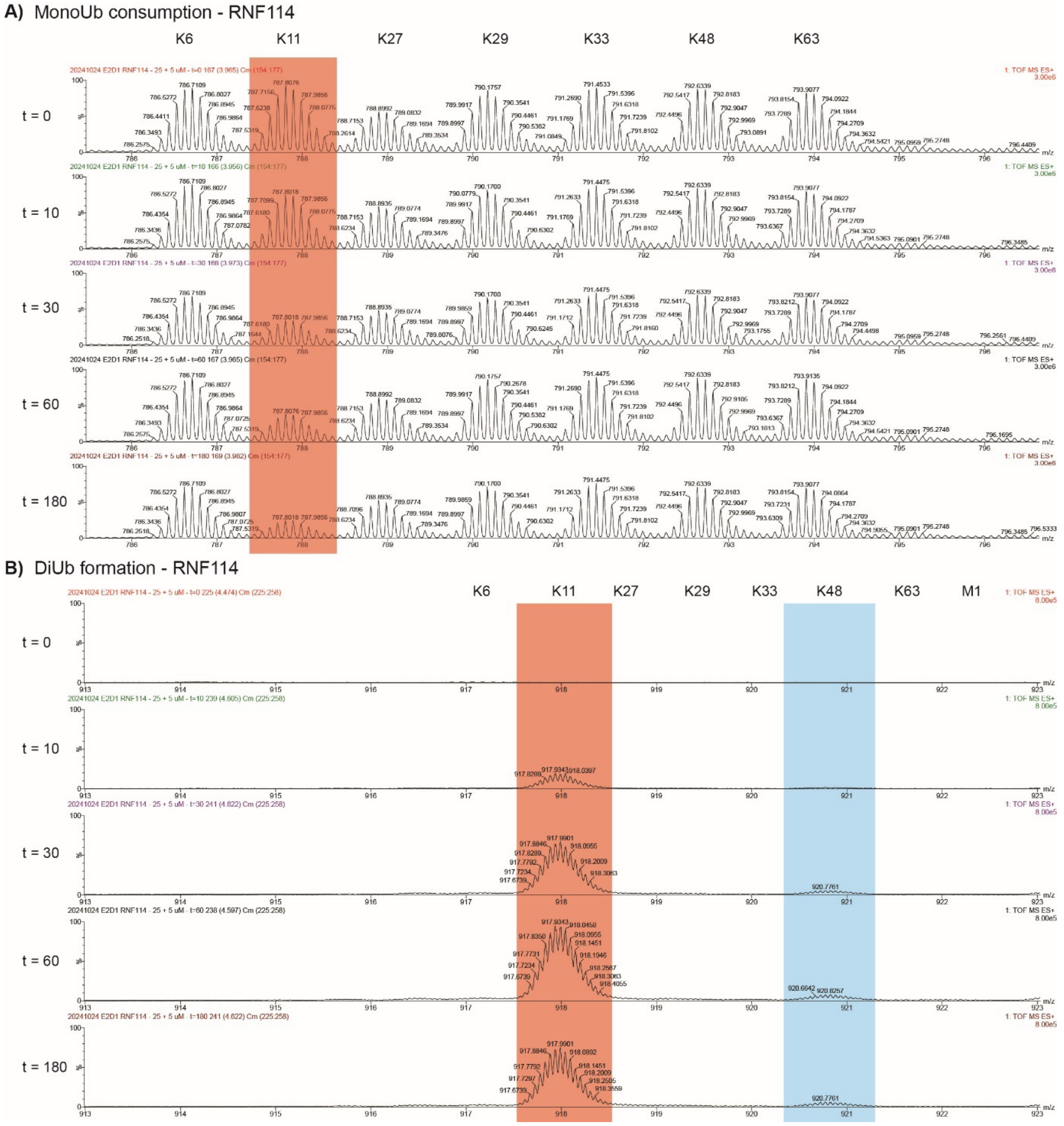


**Supplementary Fig. 6. RNF114 elongates Ub via K11 linkage.** RNF114 prefers to elongate Ub on K11, as shown by neutron encoded **A**) mono-Ub consumption and **B**) di-Ub formation,

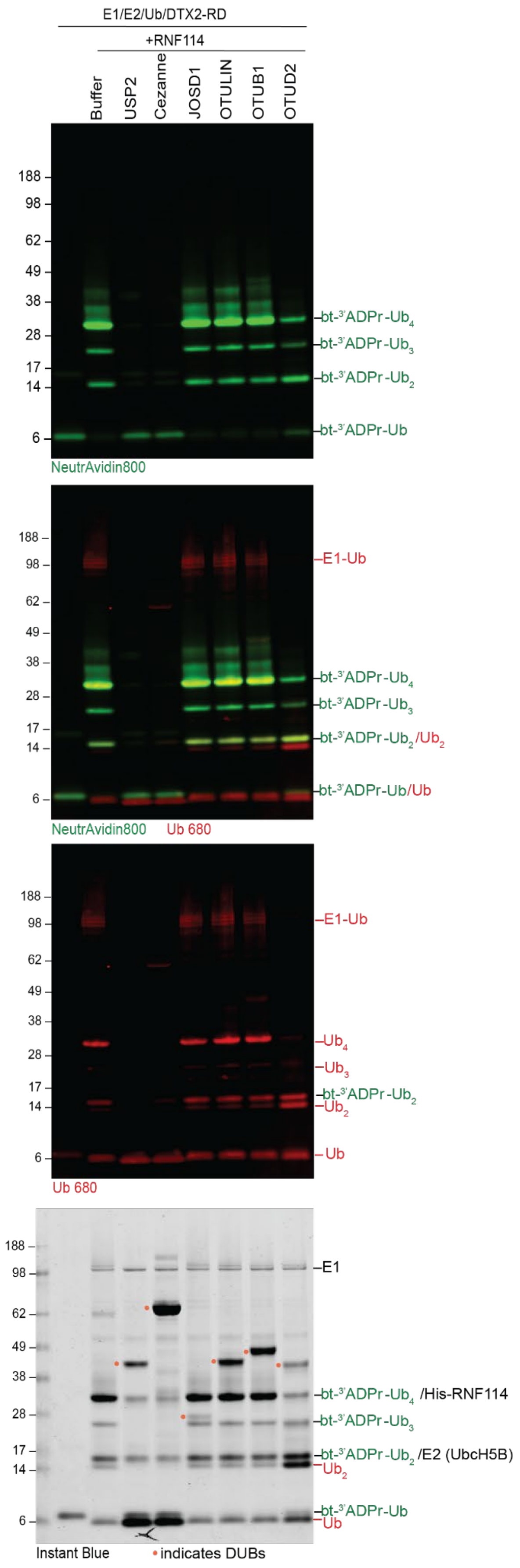


**Supplementary Fig. 7. DUB activity on the product of DTX2 and RNF114**. Panel of DUBs to remove RNF114-catalysed poly-ubiquitination of biotin-$^{3'}$ADPr-Ub probe 1.

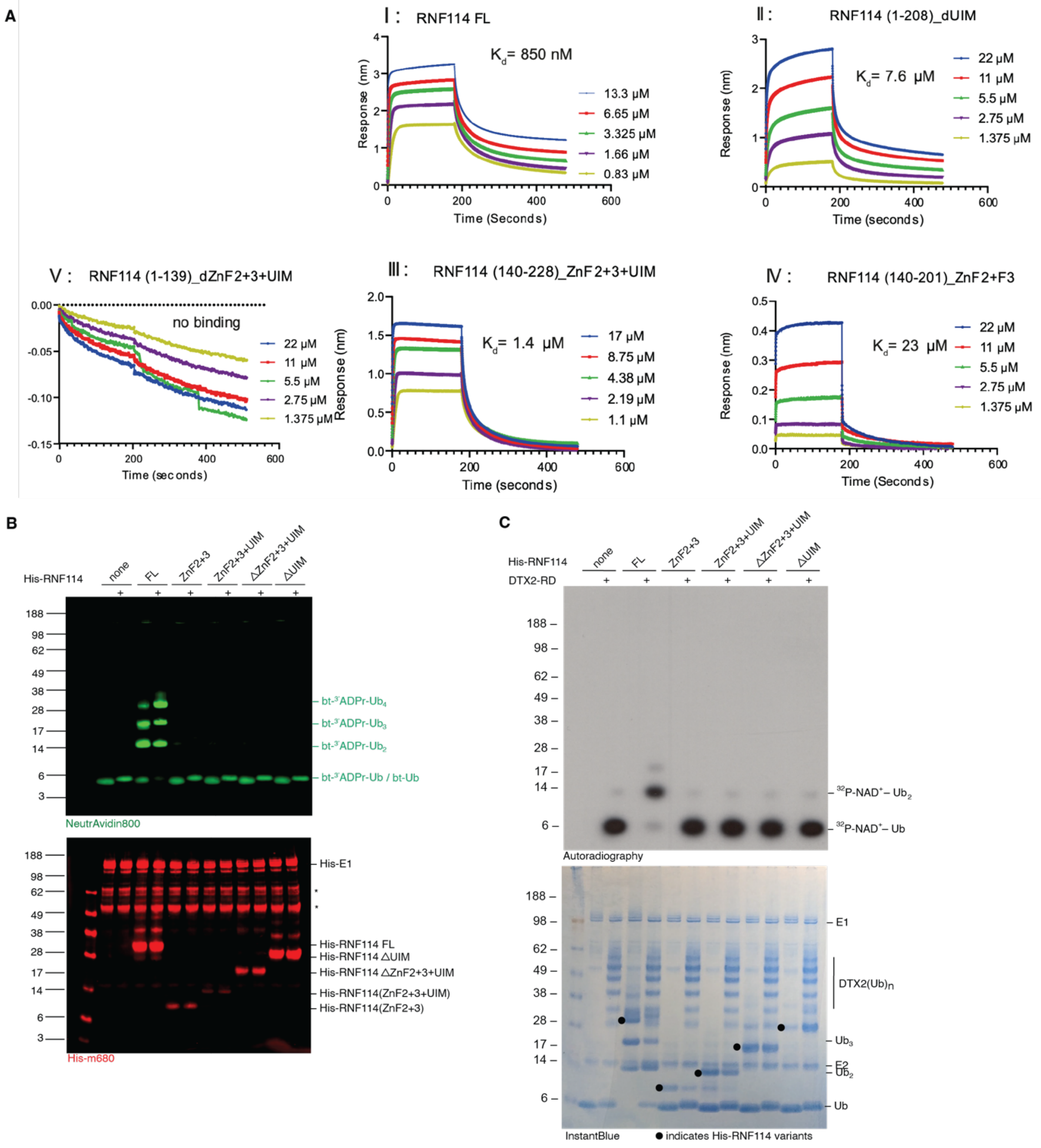


**Supplementary Fig. 8. RNF114 ZnF2+ZnF3+UIM domains are responsible for $^{3'}$ADPr-Ub binding**. A)) BLI binding curve of RNF114 FL and truncation mutants to $^{3'}$ADPr-Ub probe 1, B) Biotin-$^{3'}$ADPr-Ub probe 1 in vitro elongation reactions with RNF114 FL and truncations, asterisks (*) denote contaminants from E1 preparation, C) $^{32}$P-NAD$^+$–Ub in vitro elongation reactions with RNF114 FL and truncations.

### Supplemental information
### Chemical synthesis

#### COSY, HSQC and HMBC NMR spectra of compound 7, validating 3’-O-alkylation

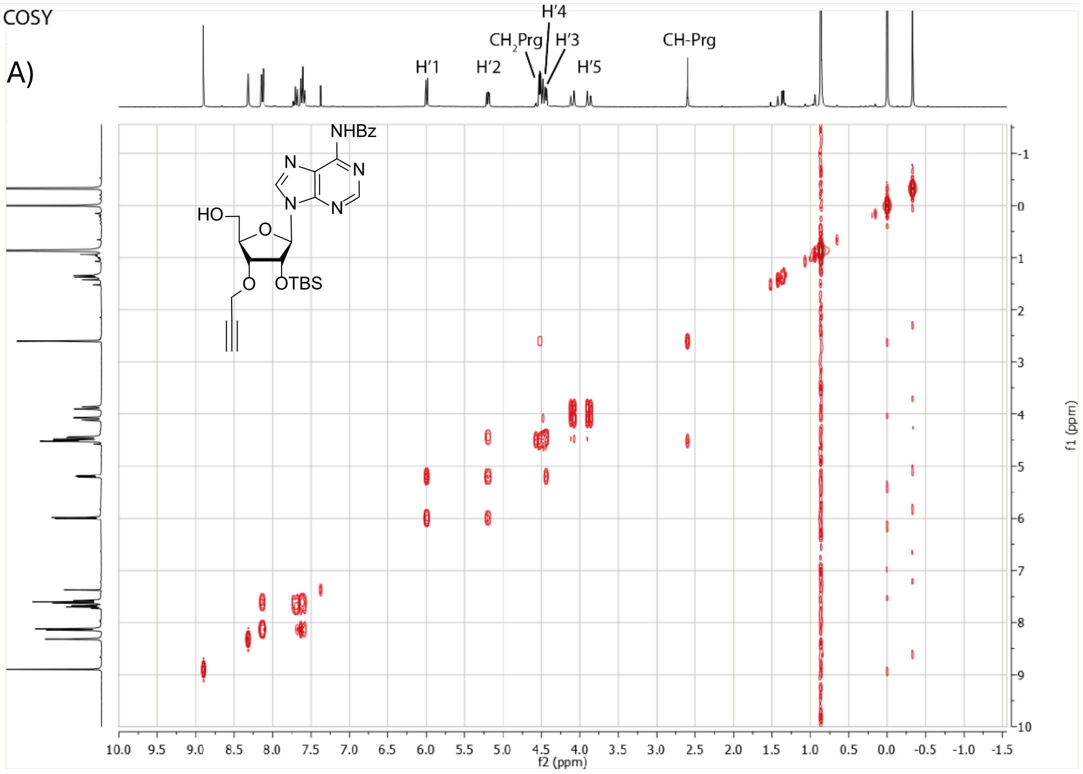


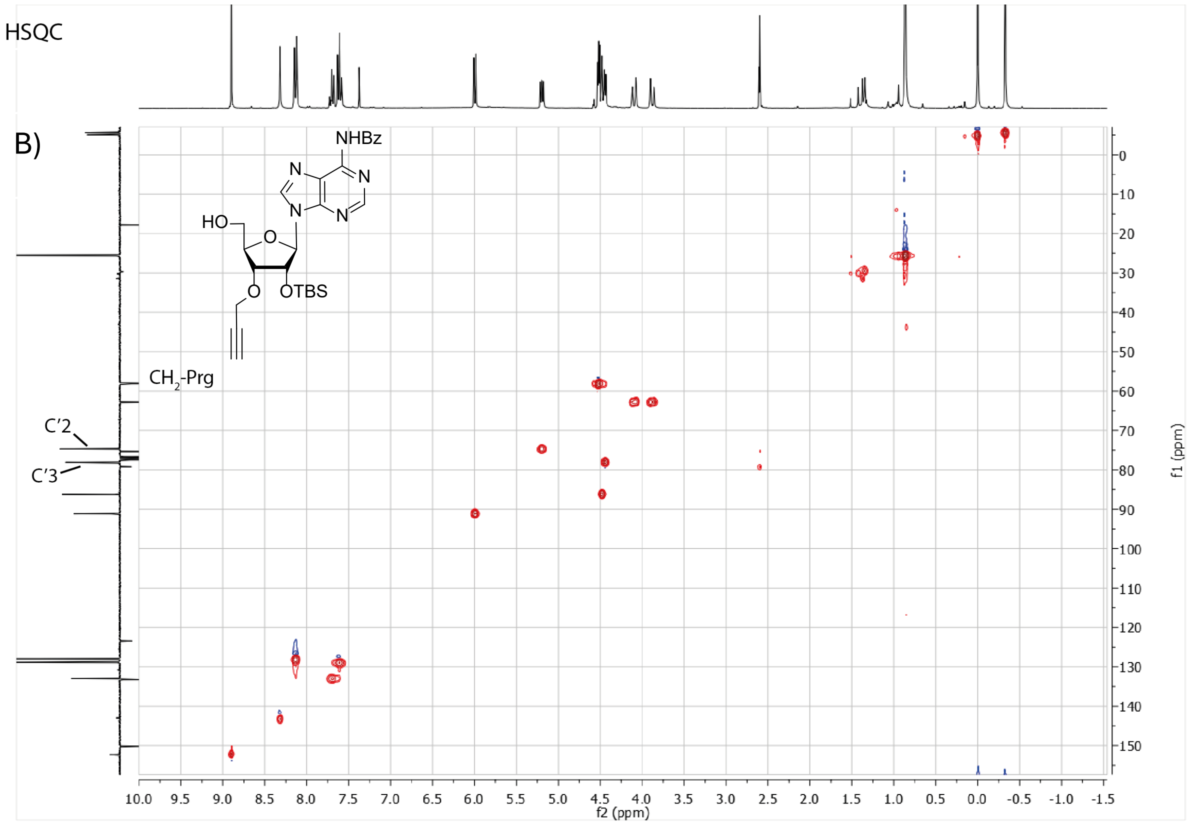

C)

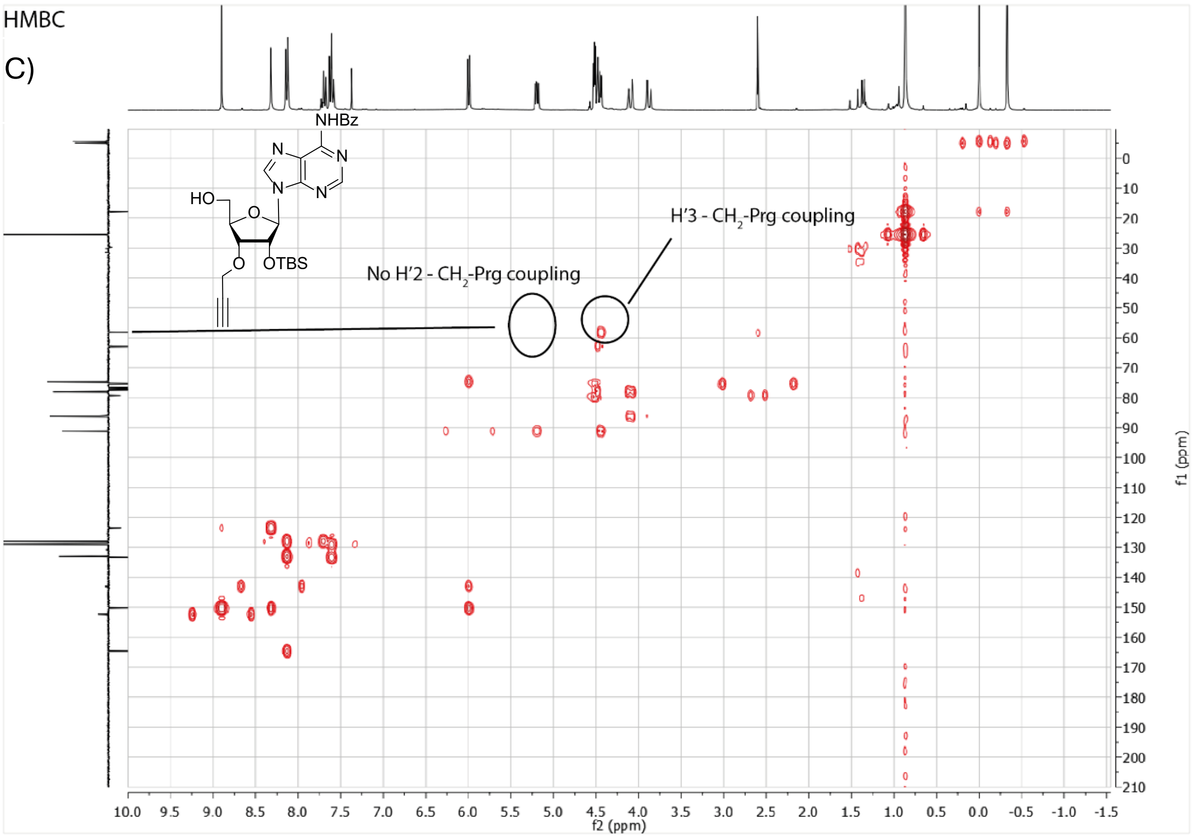


**Supplementary. Fig. 9. A)** COSY NMR spectra compound **21**. **B)** HSQC NMR spectra of **7** **C)** HMBC spectra of **7**, crucial couplings verifying 3'-*O* alkylation are annotated.

#### Chemical Synthesis

##### General synthetic procedures

All reagents were used as received unless stated otherwise. Solvents used in synthesis were dried and stored over 4Å molecular sieves, except for MeOH and MeCN which were stored over 3Å molecular sieves. Triethylamine (TEA) and diisopropylethylamine (DIPEA) were stored over KOH pellets. Column chromatography was performed on silica gel 60 Å (40-63 µm, Macherey-Nagel). TLC analysis was performed on Macherey-Nagel aluminium sheets (silica gel 60 $F_{254}$). TLC was used to visualize compounds by UV at wavelength 254 nm and by spraying with either cerium molybdate spray (25 g/L $(NH_4)_6Mo_7O_{24}$, 10 g/L $(NH_4)_4Ce(SO_4)_4 \cdot H_2O$ in 10% $H_2SO_4$ water solution) or $KMnO_4$ spray (20 g/L $KMnO_4$ and 10 g/L $K_2CO_3$ in water) followed by charring at c.a. 250 °C. NMR spectra were recorded on a Bruker AV-300 NMR. Chemical shifts (δ) are given in ppm relative to tetramethyl silane. Coupling constants (*J*) are given in Hz. All given $^{13}C$-APT spectra are proton decoupled.

##### LC-MS measurements and HPLC purifications

LC-MS measurements were conducted on a Waters ACQUITY UPLC H-class System equipped with a Waters ACQUITY Quaternary Solvent Manager (QSM), Waters ACQUITY UPLC Photodiode Array (PDA) eλ Detector (λ = 210-800 nm), Waters

ACQUITY UPLC Protein BEH C18 column (1.7 µM, 2.1 x 50 mm) and LCT Premier Orthogonal acceleration Time of Flight Mass Spectrometer (m/z = 100-1600) in ES+ mode. Samples were run for 3min at 40 °C using 2 mobile phases: A: MQ + 0.1% formic acid, B: MeCN + 0.1% formic acid. Gradient: 0 - 95% B at a flow rate of 0.5 mL/min. Data processing was performed using Waters MassLynx Mass Spectrometry Software 4.1 (deconvolution with MaxEnt1 function).
HPLC purification was performed on a **A)** Shimadzu semi-preparative RP-HPLC system, equipped with a Waters C18-Xbridge 5 µm OBD (10 x 150 mm) column at a flowrate of 6.5 mL/min. using 2 mobile phases: A: MQ + 0.05% FA, B: MeCN + 0.05 % FA. Gradient: 10 -> 70% B. HPLC system **B**) Waters preparative RP-HPLC system, equipped with a Waters C18-Xbridge 5 µm OBD (30 x 150 mm) column at a flowrate of 37.5 mL/min using 3 mobile phases: A: MQ, B: $CH_3CN$ and C: 1% TFA in MQ. Gradient: 20 -> 45% B, 5% C. High resolution mass spectra were recorded on a Waters XEVO-G2 XS Q-TOF mass spectrometer equipped with an electrospray ion source in positive mode (source voltage 3.0 kV, desolvation gas flow 900 L/hr, temperature 250 °C) with resolution R = 22000 (mass range m/z = 50-2000) and 200 pg/µL Leu-Enk (m/z = 556.2771) as a "lock mass".

**$N^6$, 2',5'-*O*-tri-trityl adenosine (3)[40,41]**

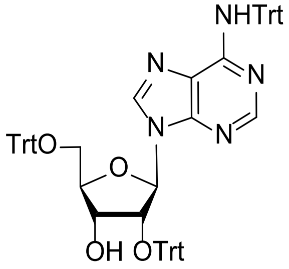


To a solution of adenosine (3.0 g, 10.9 mmol) in pyridine (160 mL) at rt was added DMAP (1.34 g, 10.9 mmol, 1 eq.) followed by trityl chloride (12.1 g, 43.4 mmol, 4 eq.). The resulting mixture was stirred at 80 °C, and after 26 h additional trityl chloride (3.07 g, 11.0 mmol, 1 eq.) was added and stirring continued at 80 °C. After 19 h, another portion of trityl chloride (4.5 g, 16.14 mmol, 1.5 eq.) was added. After another 6 h, the procedure was repeated for the final time, with the addition of trityl chloride (4.50 g, 16.15 mmol 2.5 eq.), and stirring continued another 15 h at 80 °C. The mixture was subsequently cooled to rt and quenched by addition of MeOH (30 mL). The reaction mixture was co-evaporated with toluene and concentrated *in vacuo*. The resulting crude material was purified by chromatography on silica gel (0% → 100% EtOAc in heptane) affording the title compound **3** (3.47 g, 32%) $^1$H NMR (300 MHz, $CDCl_3$) δ 7.94 (d, $J$ = 12.4 Hz, 2H), 7.34 – 7.17 (m, 45H), 6.37 (d, $J$ = 7.3 Hz, 1H), 5.17 (dd, $J$ = 7.4, 4.6 Hz, 1H), 4.09 (t, $J$ = 3.7 Hz, 1H), 3.30 (dd, $J$ = 10.5, 3.7 Hz, 1H), 3.04 (dd, $J$ = 10.5, 3.4 Hz, 1H), 2.90 (d, $J$ = 4.6 Hz, 1H). $^{13}$C NMR (75 MHz, $CDCl_3$) δ 154.1, 152.4, 149.4, 145.1, 143.5, 143.2, 139.5, 129.1, 128.7, 128.3, 128.3, 128.0, 127.9, 127.7, 127.3, 127.0, 125.4, 121.3, 87.9, 87.2, 86.5, 84.4, 77.1, 71.5, 70.7, 63.9. Spectroscopic data was in agreement with literature.[41a]

**$N^6$, 2’,5’-*O*-tri-trityl-3’-*O*-propargyl adenosine (4)**

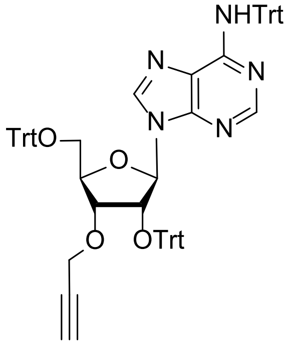

Compound **3** (1.5 g, 1.51 mmol) was co-evaporated (3x) with toluene. The resulting residue was put under argon and dissolved in anhydrous DMF (7.6 mL, 0.2 M) and cooled to 0 °C. Subsequently, NaH (0.091 g, 3.77 mmol, 2.5 eq.) was added as a solid in one portion followed by propargyl bromide (0.45 g, 3.77 mmol, 2.5 eq.), which was added dropwise. After 30 minutes, the reaction mixture was quenched with MeOH and diluted with sodium bicarbonate. Then, the aqueous layer was three times washed with diethyl ether and the combined organic layers were washed with brine. The combined organic layers were dried over $MgSO_4$, filtered, and concentrated *in vacuo*. The resulting orange-brown oil was purified by flash column chromatography on silica gel (0% → 40% EtOAc in heptane) affording the title compound **4** (1.0 g, 64.0%). $^1$H NMR (300 MHz, $CDCl_3$) δ 7.67 (s, 2H), 7.40 – 7.00 (m, 45H), 5.98 (d, *J* = 6.9 Hz, 1H), 5.31 – 5.26 (m, 1H), 4.30 – 4.24 (m, 1H), 3.96 (dd, *J* = 15.5, 2.4 Hz, 1H), 3.78 (dd, *J* = 15.5, 2.4 Hz, 1H), 3.35 – 3.27 (m, 2H), 3.13 (dd, *J* = 10.1, 4.4 Hz, 1H), 2.29 (t, *J* = 2.4 Hz, 1H). $^{13}$C NMR (75 MHz, $CDCl_3$) δ 153.8, 151.9, 148.6, 145.1, 143.6, 143.6, 140.0, 129.0, 128.9, 128.7, 127.9, 127.7, 127.7, 127.3, 127.1, 126.9, 87.7, 87.4, 86.9, 82.2, 79.7, 78.1, 74.6, 74.0, 71.3, 63.2, 57.7.

**2’,5’-di-*O*-*tert*-butyldimethylsilyl-3’-*O*-propargyl adenosine (5)**

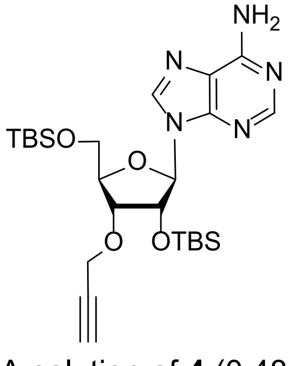

A solution of **4** (0.48 g, 0.90 mmol) in 80% AcOH:$H_2O$ (5 mL) was stirred at 80 °C for 16 h. The reaction mixture was then concentrated *in vacuo* and co-evaporated with toluene, affording the crude 3’-*O*-propargyl ribose, LC-MS (M + $H^+$ = 306.077). Subsequently, the crude was dissolved in anhydrous DMF (3.22 mL, 0.2 M) and placed under argon before imidazole (0.16 g, 2.41 mmol, 3.75 eq.) and TBS-Cl (0.29 g, 1.93 mmol, 3 eq.) were added. After stirring for 17 h at 50 °C, the reaction was quenched with MeOH. The mixture was transferred to a separatory funnel and $Et_2O$ (25 mL) and 1 M HCl (5 mL) were added. The layers were separated, and the aqueous layer was extracted with $Et_2O$. The combined organic layers were washed with brine. The organic layer was dried over $MgSO_4$, filtered, and concentrated *in vacuo* to obtain a yellow oil. The yellow oil was purified by flash column chromatography on silica gel (0% → 45% EtOAc in heptane) affording the title compound **5** (0.249 g, 0.47 mmol, 52%).$^1$H NMR (300 MHz, $CDCl_3$) δ 8.18 (s, 1H), 8.14 (s, 1H), 6.19 (s, 1H), 5.92 (d, *J* = 4.8 Hz, 1H), 4.55 (t, *J* = 4.8, 3.9 Hz, 1H), 4.17 (dd, *J* = 4.5, 2.4 Hz, 2H), 4.14 – 4.09 (m, 2H), 3.79 (ddd, *J* = 55.6, 11.5, 2.5 Hz, 2H), 2.28 (t, *J* = 2.4 Hz, 1H), 0.82 (s, 9H), 0.68 (s, 9H), 0.01 (d, *J* = 4.3 Hz, 6H), -0.13 (s, 3H), -0.26 (s, 3H). $^{13}$C NMR (75 MHz, $CDCl_3$) δ 154.6, 151.3, 149.7, 139.6, 119.8, 88.6, 88.6, 83.2, 83.2, 79.2, 76.2, 75.7, 75.1, 62.5, 57.7, 29.7, 26.0, 25.5, -5.1, -5.1, -5.3, -5.4.

**$N^6$-benzoyl-2',5'-di-*O*-*tert*-butyldimethylsilyl-3'-*O*-propargyl adenosine (6)**

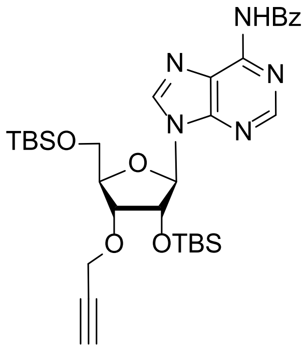


Compound **5** (0.23 g, 0.43 mmol) dissolved in anhydrous pyridine (1.88 mL, 0.2 M) was placed under argon and cooled to 0 °C. Next, benzoyl-Cl (0.16 mL, 1.55 mmol, 3.6 eq.) was added dropwise. After stirring for 18 h, $NH_3OH$ in MeOH (1.88 mL, 1.5 M) and THF (1.88 mL) were added (Pyr: $NH_3$ in MeOH: THF 1:1:1 v/v/v) and allowed to react for 45 min. The reaction mixture was concentrated *in vacuo* and co-evaporated with toluene to obtain a pale white solid. The solid was redissolved and purified by flash column chromatography (0% → 60% EtOAc in heptane) affording the title compound **6** (0.250 g, 0.37 mmol, 87%). $^1$H NMR (300 MHz, $CDCl_3$) δ 8.79 (s, 1H), 8.44 (s, 1H), 8.10 – 7.97 (m, 2H), 7.62 – 7.46 (m, 3H), 6.15 (d, *J* = 4.8 Hz, 1H), 4.73 (t, *J* = 4.8, 4.0 Hz, 1H), 4.32 (dd, *J* = 4.5, 2.4 Hz, 2H), 4.30 – 4.26 (m, 2H), 3.94 (ddd, *J* = 54.0, 11.5, 2.5 Hz, 2H), 2.43 (t, *J* = 2.4 Hz, 1H), 0.96 (s, 9H), 0.82 (s, 9H), 0.16 (s, 3H), 0.14 (s, 3H), 0.02 (s, 3H), -0.13 (s, 3H). $^{13}$C NMR (75 MHz, $CDCl_3$) δ 164.9, 152.6, 151.7, 149.6, 141.5, 133.8, 132.8, 128.9, 128.1, 123.0, 88.8, 83.4, 79.2, 77.6, 77.2, 76.7, 76.2, 75.8, 75.4, 62.6, 57.8, 32.0, 26.2, 25.7, -4.9, -4.9, -5.2, -5.3.

**N^6^-benzoyl-2'-*O*-*tert*-butyldimethylsilyl-3'-*O*-propargyl adenosine (7)**

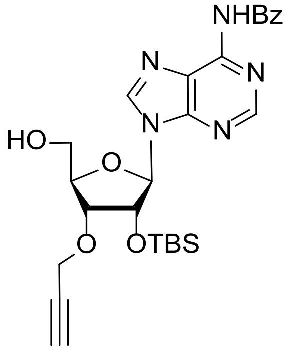


To an ice-cooled solution of **6** (0.25 g, 0.39 mmol) in THF (3.9 mL, 0.1 M), a freshly prepared TFA:$H_2O$ mixture (3 mL, 1:1, v/v, 50 eq.) was gradually added. After stirring for 4 h at 0 °C, the solution was quenched by the careful addition of solid $NaHCO_3$ until the pH reached 7. The solution was further diluted with sat. aq. $NaHCO_3$ (40 mL) and extracted three times with EtOAc (3 x 30 mL). The resulting organic layers were combined, dried over $MgSO_4$, filtered and concentrated *in vacuo*. Flash column chromatography (0% -> 100% EtOAc in heptane) afforded the titled compound **7** as a white solid (0.13 g, 0.25 mmol, 64%).$^1$H NMR (300 MHz, $CDCl_3$) δ 8.79 (s, 1H), 8.20 (s, 1H), 8.04 – 7.99 (m, 2H), 7.62 – 7.46 (m, 3H), 5.88 (d, *J* = 7.3 Hz, 1H), 5.08 (dd, *J* = 7.3, 4.7 Hz, 1H), 4.40 (dd, *J* = 4.3, 2.4 Hz, 2H), 4.37 – 4.31 (m, 2H), 3.87 (ddd, *J* = 64.6, 13.0, 1.7 Hz, 2H), 2.49 (t, *J* = 2.4 Hz, 1H), 0.75 (s, 9H), -0.11 (s, 3H), -0.44 (s, 3H). $^{13}$C NMR (75 MHz, $CDCl_3$) δ 164.7, 152.4, 150.4, 150.3, 143.0, 133.4, 133.1, 129.0, 128.1, 123.5, 91.2, 86.3, 79.3, 78.2, 75.4, 74.8, 62.9, 29.8, 25.6, -5.0, -5.6.

**5'-*O*-(*N*^6^-benzoyl-2'-*O*-*tert*-butyldimethylsilyl-3'-*O*-propargyl)-2-cyanoethyl-*N*-*N*-diisopropylphosphoramidite (8)**

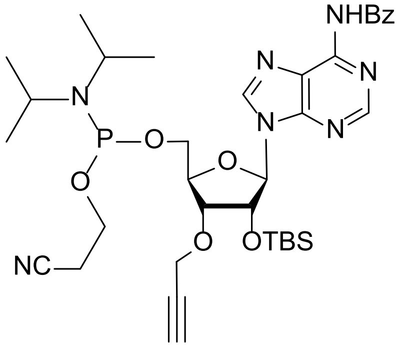


Compound **7** (99 mg, 0.189 mmol) was co-evaporated three times with toluene and dissolved in anhydrous DCM (0.96 mL, 0.2 M) under argon atmosphere. DIPEA (0.1 mL, 0.57 mmol, 3.0 eq.) and 2-cyanoethyl-N,N-diisopropylchlorophosphoramidite (0.11 mL, 0.48 mmol, 2.5 eq.) were added to the reaction mixture. After 2 hours, TLC indicated full conversion. Subsequently, the reaction mixture was diluted in DCM and flash column chromatography (0 % → 60% EtOAc in heptane + 1% TEA) afforded titled compound **8** as a white solid (0.11 g, 0.15 mmol, 79%). $^1$H NMR (300 MHz, $CDCl_3$) δ 9.20 (s, 1H), 8.86 (s, 1H), 8.51 (d, *J* = 6.1 Hz, 1H), 8.14 – 8.02 (m, 2H), 7.72 – 7.52 (m, 3H), 6.19 (dd, *J* = 8.3, 4.5 Hz, 1H), 4.90 (dt, *J* = 16.3, 4.4 Hz, 1H), 4.48 – 4.42 (m, 2H), 4.41 – 4.31 (m, 2H),

4.17 (q, $J$ = 7.1 Hz, 1H), 4.09 – 3.88 (m, 2H), 3.95 – 3.80 (m, 2H), 3.76 – 3.38 (m, 3H), 2.90 – 2.79 (m, 2H), 2.79 – 2.53 (m, 2H), 2.51 (q, $J$ = 2.3 Hz, 1H), 1.33 – 1.24 (m, 12H), 0.90 (d, $J$ = 7.2 Hz, 9H), 0.10 (d, $J$ = 9.5 Hz, 3H), -0.05 (d, 3H). $^{13}$C NMR (75 MHz, $CDCl_3$) δ 164.6, 152.7, 149.4, 139.2, 133.8, 132.7, 128.9, 127.8, 117.6, 89.4, 88.8, 82.2, 82.1, 82.0, 81.9, 79.3, 77.5, 77.0, 76.6, 76.3, 75.2, 74.9, 62.0, 61.8, 58.8, 58.5, 57.9, 57.8, 43.3, 43.2, 43.1, 43.0, 29.7, 25.6, 25.6, 24.8, 24.8, 24.7, 24.7, 20.5, 20.4, 18.0, 17.9, -5.0. $^{31}$P NMR (121 MHz, $CD_3CN$-$d_3$) δ 148.82, 148.79.

**1’-*O*-methyl-2’,3’-*O*-di-benzoyl-5-(*O*-di-fluorenylmethyl phosphate) ribofuranoside (9)**

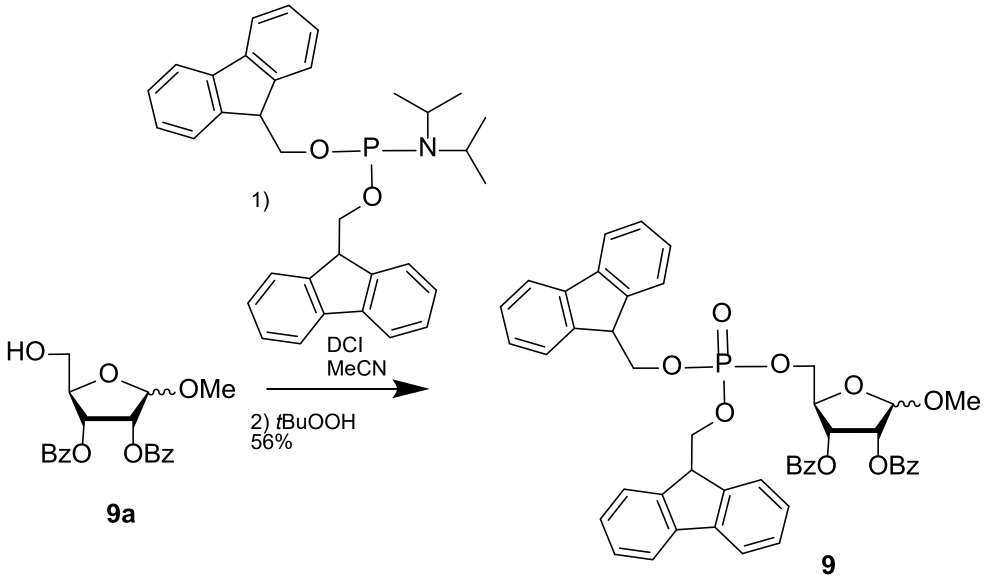


Compound **9a** (0.11 g, 0.30 mmol) was synthesized as reported previously[42] and subsequently co-evaporated with toluene and put under argon. Subsequently, a 0.2M stock solution of Bis-(9H-fluoren-9-ylmethyl)-N,N-diisopropylamidophosphoramidite (0.17 g, 0.33 mmol, 1.1 eq.) in anhydrous MeCN (1.5 mL) was added. The solution was cooled to 0 °C and DCI (85 mg, 0.72 mmol, 2.4 eq.) was added as a solution in MeCN (3.5 mL, 0.2 M). After stirring at 0 °C for 15 min, the mixture was allowed to warm to rt following stirring for an additional 2 h. The mixture was cooled to 0°C and *t*BuOOH (0.55 mL, 3.0 mmol, 10 eq.) as a 5.5 M solution in nonane was added dropwise. The mixture was stirred for an additional 1.5 h and quenched with $H_2O$. Next, the reaction mixture was extracted three times with DCM. The combined organic layers were dried over $MgSO_4$, filtered, and concentrated *in vacuo*. Flash column chromatography (0 % -> 45% EtOAc in heptane) afforded titled compound **9** (0.13 g, 0.16 mmol, 52%). $^{1}$H NMR (300 MHz, $CDCl_3$) δ 8.03 – 7.98 (m, 2H), 7.90 – 7.85 (m, 2H), 7.74 – 7.68 (m, 4H), 7.61 – 7.50 (m, 6H), 7.44 – 7.37 (m, 4H), 7.36 – 7.26 (m, 8H), 5.67 – 5.62 (m, 1H), 5.60 (d, $J$ = 5.7 Hz, 1H), 5.11 (s, 1H), 4.56 – 4.50 (m, 1H), 4.36 – 4.29 (m, 4H), 4.23 – 4.10 (m, 4H), 3.35 (s, 3H). $^{13}$C NMR (75 MHz, $CDCl_3$) δ 165.3, 165.2, 143.1, 143.1, 141.4, 133.5, 133.4, 129.8, 129.7, 129.2, 128.9, 128.5, 128.4, 127.9, 127.1, 125.2, 120.0, 106.5, 79.7, 77.5, 77.1, 76.7, 75.3, 72.1, 69.5, 69.4, 68.1, 55.4, 47.9, 47.9. $^{31}$P NMR (121 MHz, Chloroform-$d$) δ -1.75.

**(5’-*O*-diphosphate-1’-*O*-methyl-riboside)-3’-*O*-propargyl-2’-*O*-*tert*-butyldimethylsilyl- adenosine (10)**

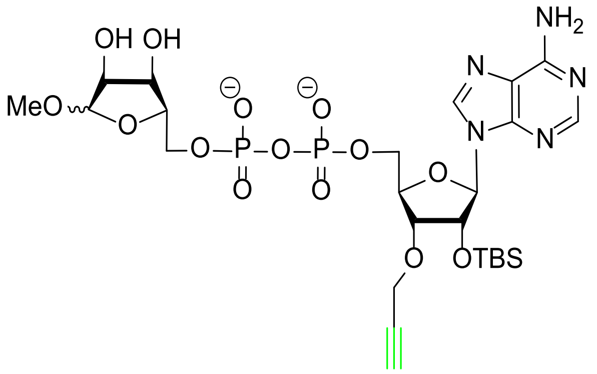


Phosphate **9** (42.6 mg, 44 µmol, 1.5 eq.) was dissolved in MeCN (0.44 mL, 0.1M) before treated with triethylamine (92 µL, 0.66 mmol, 15 eq.). Upon full conversion, monitored by LC-MS (**8**, M + $H^+$ = 453) the reaction mixture was concentrated *in vacuo* and co-evaporated with pyridine to form the pyridinium salt, which was subsequently co-evaporated with toluene and placed under argon atmosphere. Meanwhile amidite **8** (20.9 mg, 29 µmol, 1 eq.) was co-evaporated with anhydrous toluene (3x), placed under argon atmosphere and dissolved in anhydrous MeCN (0.48 mL). The amidite solution was added to the crude phosphate pyridinium salt followed by the addition of DCI (8.25 mg, 72 µmol, 2.5 eq.) to start the reaction. After 30 min of stirring at rt, *t*BuOOH (52 µL, 290 µmol, 10 eq.), as a 5.5 M solution in nonane was added and stirring continued for another 30 min. Next, DBU (28 µL, 187 µmol, 6.5 eq.) was added to the reaction mixture and successful formation of the pyrophosphate bridge was verified by LC-MS (M + $H^+$ = 1036). Deprotection of the benzoyl groups was facilitated by the addition of sat. aq. $NH_3OH$ (1.5 mL) and dioxane (0.5 mL), and stirred overnight. The reaction mixture was diluted with toluene and concentrated *in vacuo* before co-evaporated another time with toluene. The crude was purified by RP-HPLC (10%-60% MeCN in $H_2O$) and pure fractions lyophilized to obtain 3’-*O*-propargylated ADPr **10** (6.2 mg, 8.5 µmol, 29%). $^1$H NMR (300 MHz, $D_2O$) δ 8.71 (s, 1H), 8.45 (s, 1H), 6.12 (d, *J* = 6.8 Hz, 1H), 4.89 – 4.87 (m, 1H), 4.85 – 4.82 (m, 1H) 4.59 (s, 1H), 4.46 – 4.39 (m, 3H), 4.32 – 4.22 (m, 3H), 4.18 – 4.09 (m, 2H), 4.05 – 3.95 (m, 2H), 3.37 (s, 3H), 2.95 (t, *J* = 2.4 Hz, 1H), 0.72 (s, 9H), -0.02 (s, 3H), -0.16 (s, 3H). $^{31}$P NMR (121 MHz, $D_2O$) δ -11.42.

**Biotin-(PEG$_2$)$_2$-Ub$_{74}$-azidoethane (11)**

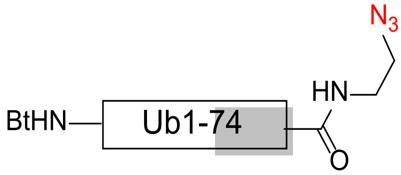


10 µmol Ub$_{1-74}$ (Arg42 → azido homoalanine) on resin[43] was treated with Bt-(PEG$_2$)$_2$-COOH (26.8 mg, 50 µmol, 5 eq.) HOBt (6.7 mg, 50 µmol, 5 eq.) and HBTU (19 mg, 50 µmol, 5 eq.) in DMF (2 mL). After 5 min of shaking, DIPEA (26 µL, 150 µmol, 15 eq.) was added. The reaction mixture was shaken overnight, after which a test cleavage confirmed

installation of the biotin LC-MS (M + $H^+$ = 8948, deconvoluted). The resin was washed with DMF and DCM. HFIP (1.5 mL, 20% in DCM) was added and the resin was shaken for 30 min at rt before filtration into a round bottom flask. The treatment was repeated once before the combined filtrate was concentrated *in vacuo* and co-evaporated with DCE three times. Subsequently, the crude ubiquitin was dissolved in a mixture of TFE:$CHCl_3$ (1.2 mL, 3:1 v/v) followed by the addition of 2-azidoethan-1-amine (25 µL, 331 µmol, 31 eq.) prepared via literature precedence[21,22], EDC-HCl (7.8 mg, 50 µmol, 5 eq.) and HOBt (6.8 mg, 50 µmol, 5 eq.). After 1 h of stirring at rt, LC-MS indicated full conversion into an azide-modified Ub (M + $H^+$ = 9018, deconvoluted). Subsequently, the mixture was concentrated *in vacuo* and co-evaporated with DCM two times. The residue was treated with TFA/TIS/$H_2O$/Phenol (90.5/2/5/2.5, v/v) for 2.5 hours before being added to an ice-cold solution of $Et_2O$:pentane (1:1, v/v). The precipitate formed was centrifuged (5 min, 3500 rpm) and the supernatant decanted. The pellet was subsequently dried with $N_2$, taken up in warm DMSO (1 mL) and diluted into warm water before purification by RP-HPLC. Pure fractions were pooled and lyophilized affording ubiquitin azide **11** (5.95 mg, 0.66 µmol, 6.6%) as a white powder. LC-MS (M + $H^+$ = 9018, deconvoluted).

**Biotin-($PEG_2$)$_2$-$Ub_{74}$-triazole-3'-*O*-ADPr (1)**

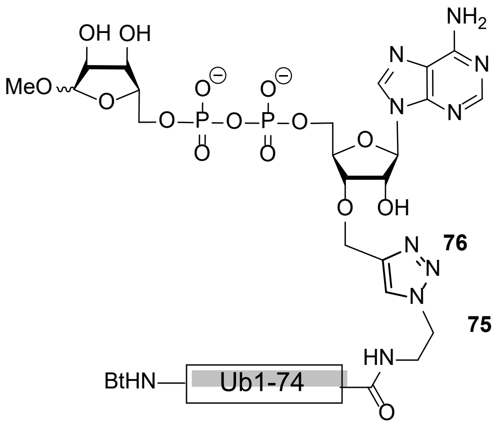


3'-*O*-propargylated ADPr **10** (183 µg, 0.3 µmol) was dissolved in MeCN (30 µL) before the addition of TBAF (0.36 µL, 0.36 µmol, 1M in THF, 1.2 eq.). After 45 min LC-MS verified removal of the TBS protecting group in **10** (M + $H^+$ = 612). Subsequently, biotin-ubiquitin$_{1-74}$-ethane-azide **11** (1 mg, 0.11 µmol) was dissolved in DMSO (30 µL). The ubiquitin solution was then added to 800 µL buffer (20 mM TRIS, 150 mM NaCl, pH 7.6) before the addition of crude 3'-*O* propargyl ADPr **10** (20 µL, 10.3 mM in MeCN, 0.21 µmol, 1.87 eq.). The mixture was then treated with 15 µL of freshly prepared click-mixture (1:1:1 v/v/v, $CuSO_4$ (100 mM in $H_2O$): Sodium Ascorbate (600 mM in $H_2O$): TBTA ligand (100 mM in MeCN) to start the reaction. The reaction mixture was shaken at 25 °C for 40 min. At this time point LC-MS indicated moderate conversion, hence 9 µL of freshly prepared click-mixture and 10 µL 3'-O propargyl ADPr **10** (0.10 µmol, 10.3 mM in MeCN, 0.94 eq.) were added. After 1 hour of total reaction time LC-MS verified complete conversion to the $^{3'}$ADPr-Ub conjugate (deconvoluted mass found: (M + $H^+$ = 9625)). The conjugate was purified by RP-HPLC and the pure fractions were pooled and lyophilized obtaining ubiquitin-ADPr conjugate **1** (0.5 mg, 0.052 μmol, 47%) as a white powder. HRMS: $C_{418}H_{692}N_{116}O_{137}P_2S$ + 7H]$^{7+}$ found: 1376.4583, calculated: 1376.5457. $C_{418}H_{692}N_{116}O_{137}P_2S$ + 8H]$^{8+}$ found: 1204.3870, calculated: 1204.6025. $C_{418}H_{692}N_{116}O_{137}P_2S$ + 9H]$^{9+}$ found: 1070.6275, calculated: 1070.8689. $C_{418}H_{692}N_{116}O_{137}P_2S$ + 10H]$^{10+}$ found: 963.6327, calculated: 963.8820. $C_{418}H_{692}N_{116}O_{137}P_2S$ + 11H]$^{11+}$ found: 876.0592, calculated: 876.3473.

#### HRMS spectra of probe 1

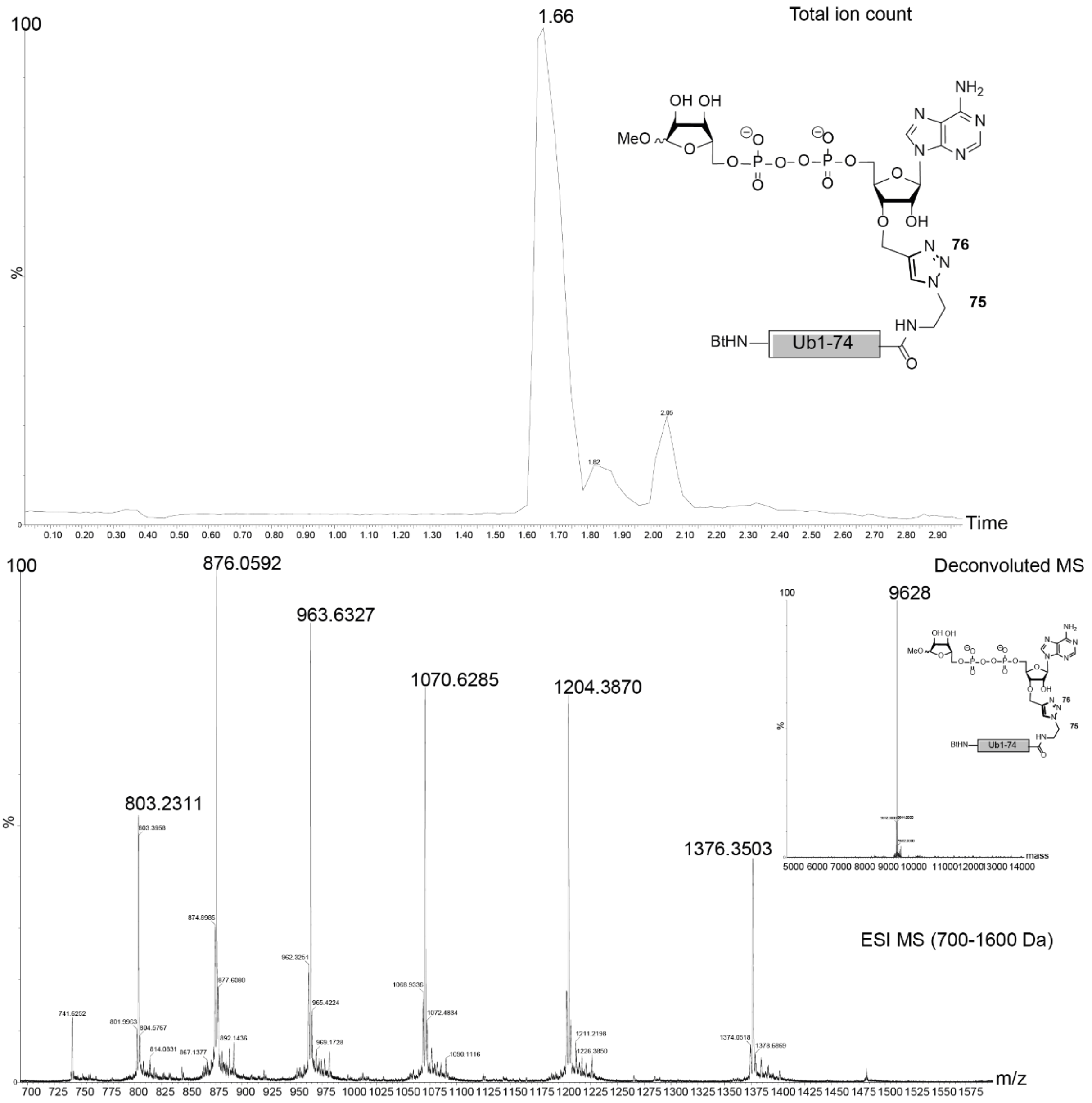


**Supplementary Fig. 10.** HRMS spectra of Biotin-$(PEG_2)_2$-$Ub_{74}$-triazole-3’-*O*-ADPr (**1**)

### NMR data

#### *3’-O-Prg-amidite and intermediates*

$^{1}H$ NMR (**3**)

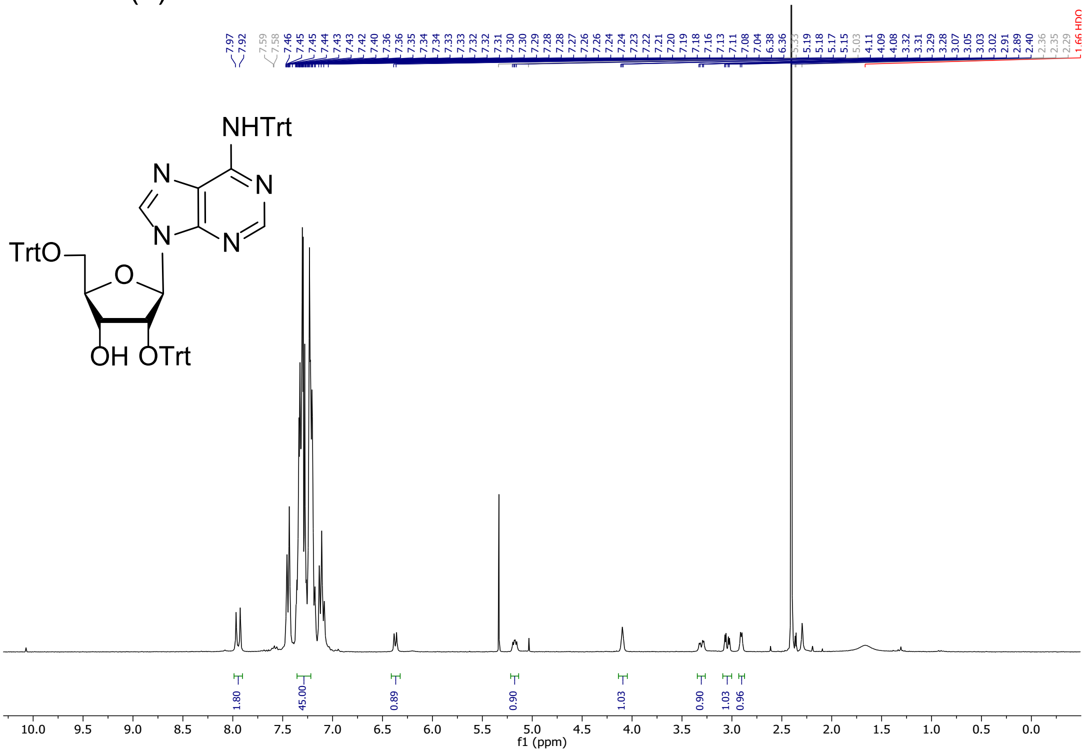


$^{13}C$ NMR (**3**)

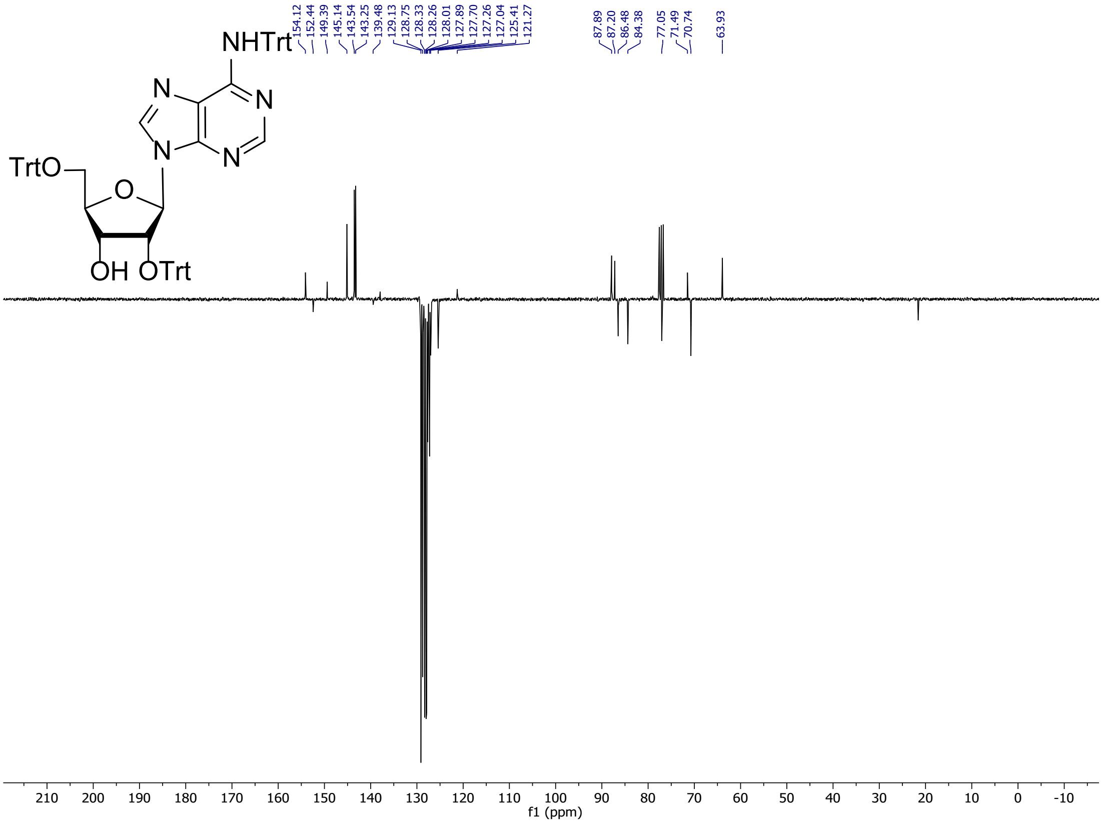

$^{1}H$ NMR (**4**)

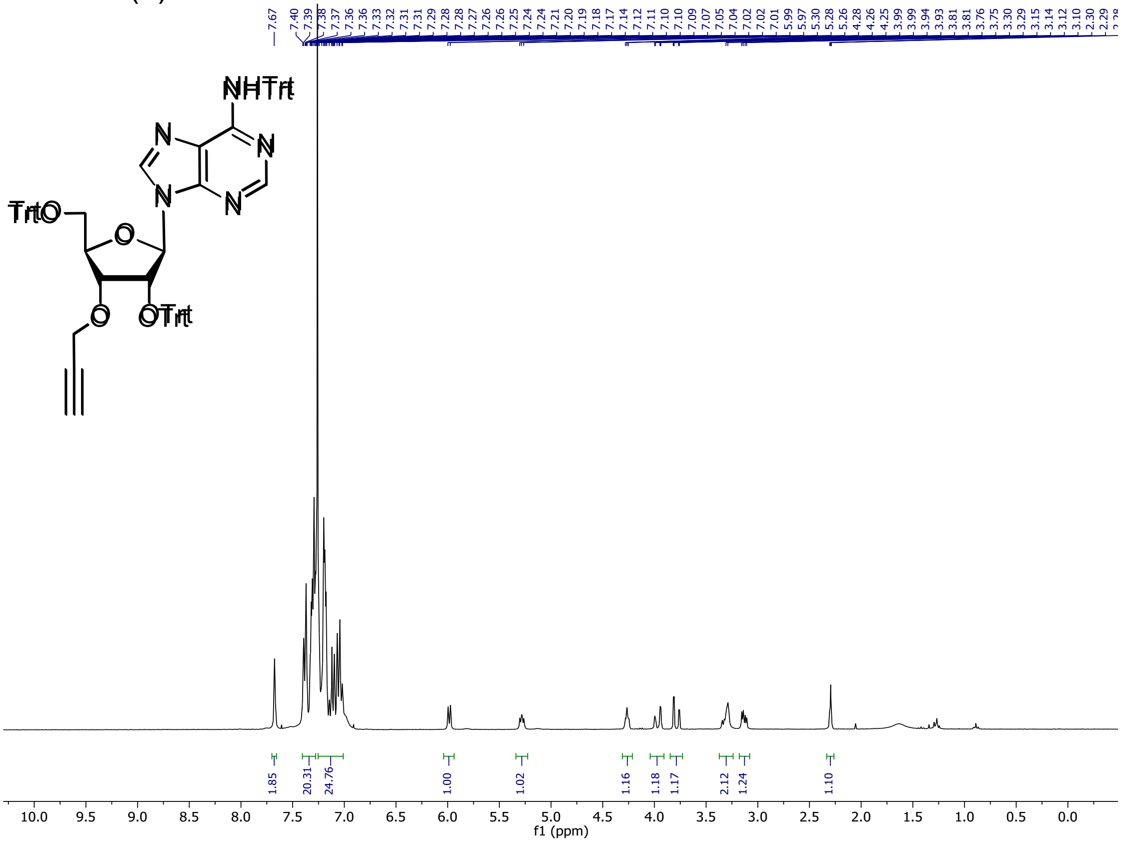

$^{13}C$ NMR (**4**)

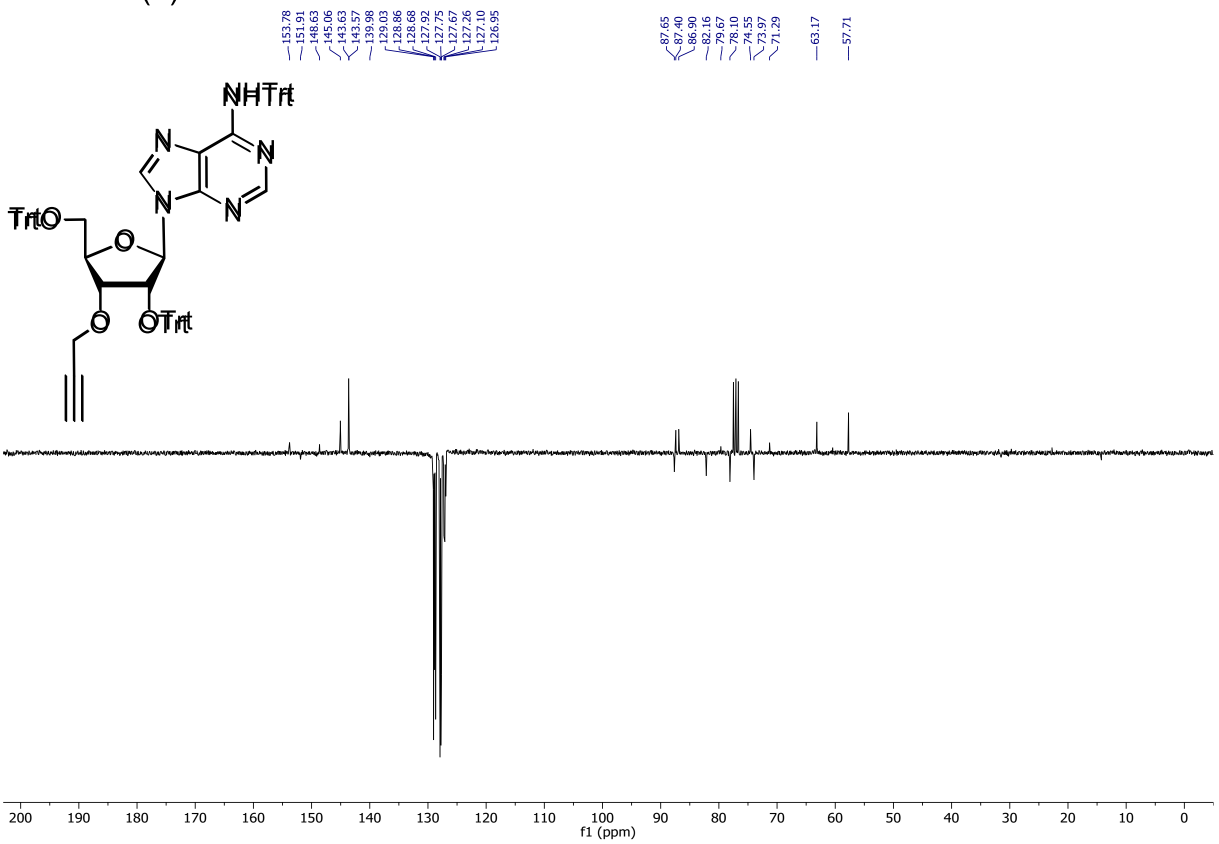

#### $^{1}$H NMR (**5**)

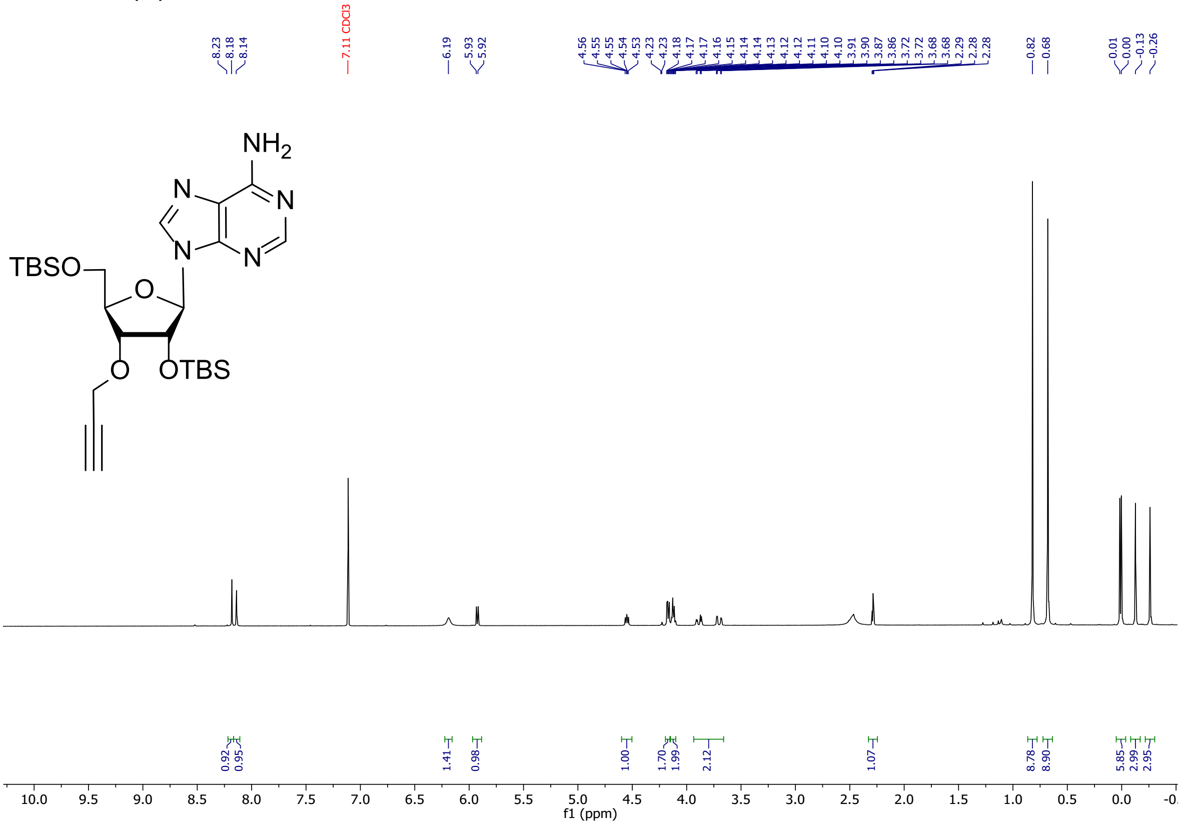


#### $^{13}$C NMR (**5**)

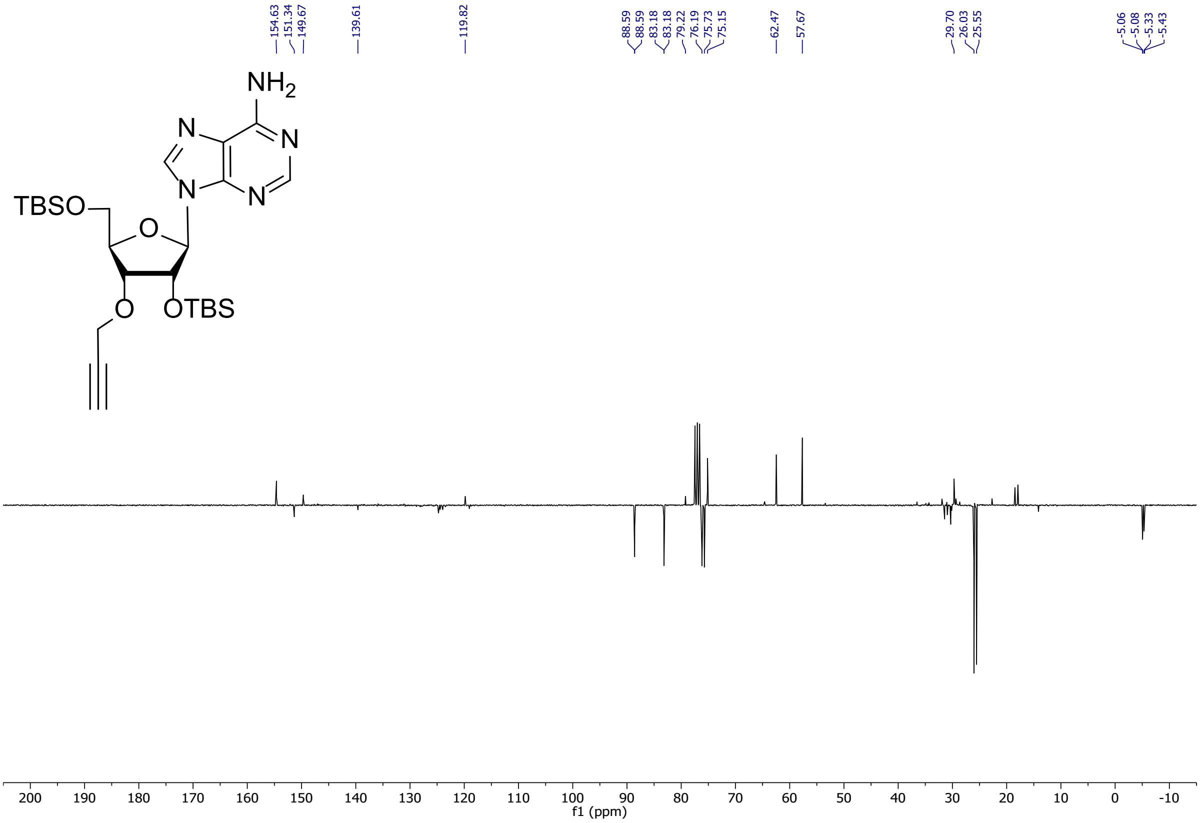

$^{1}H$ NMR (**6**)

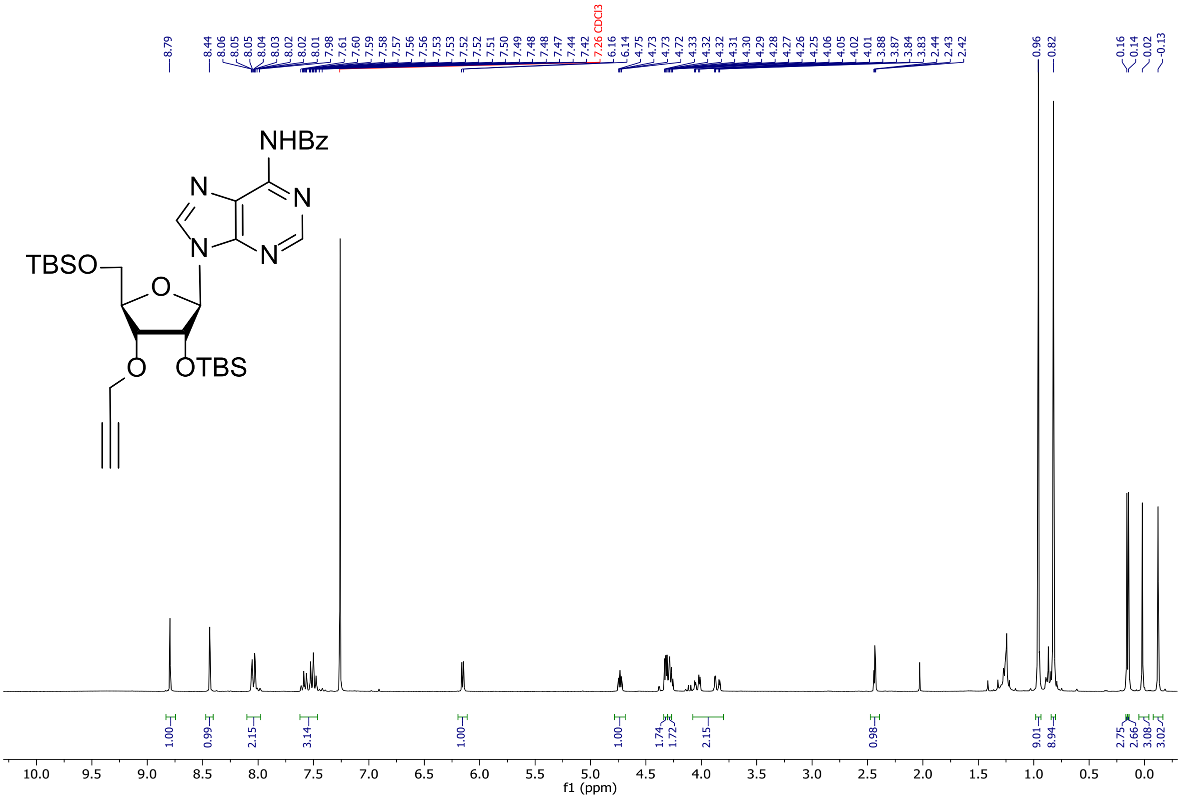

$^{13}C$ NMR (**6**)

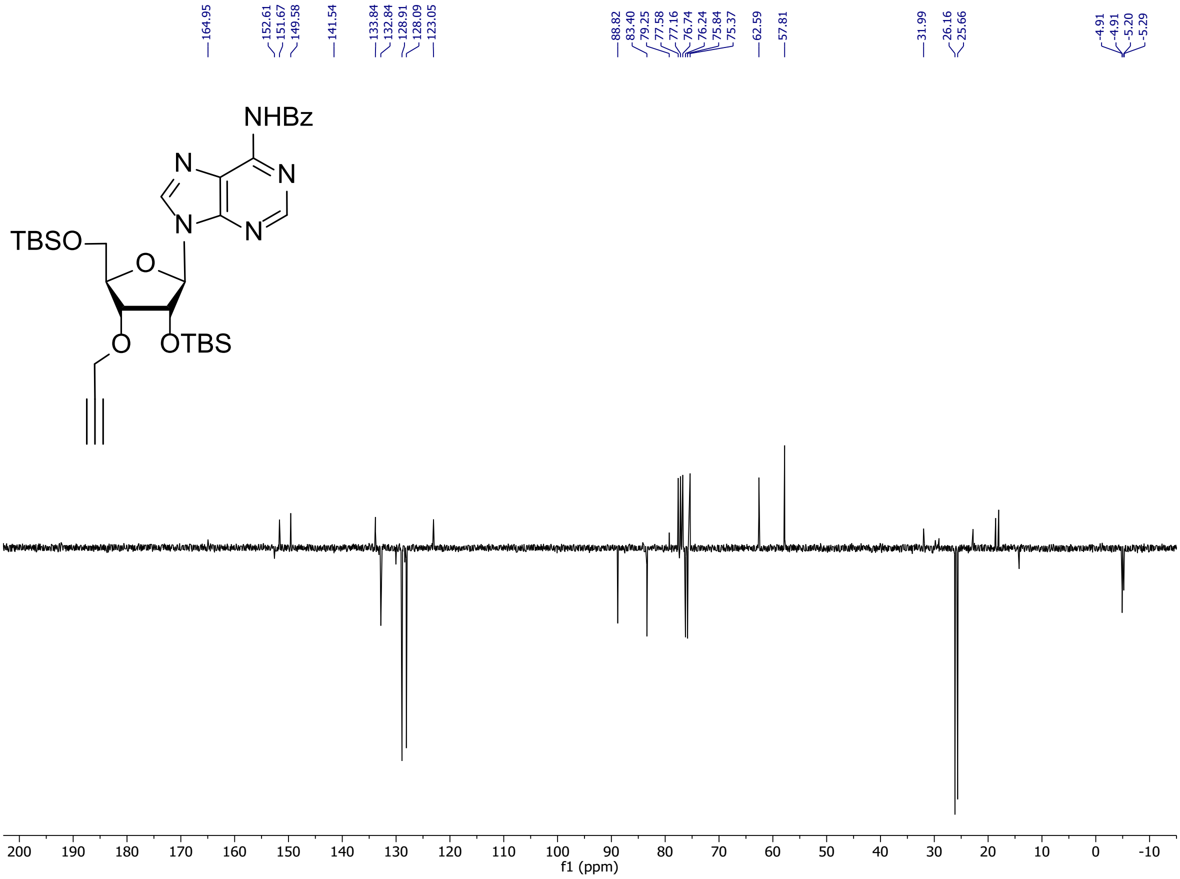

#### $^{1}H$ NMR (7)

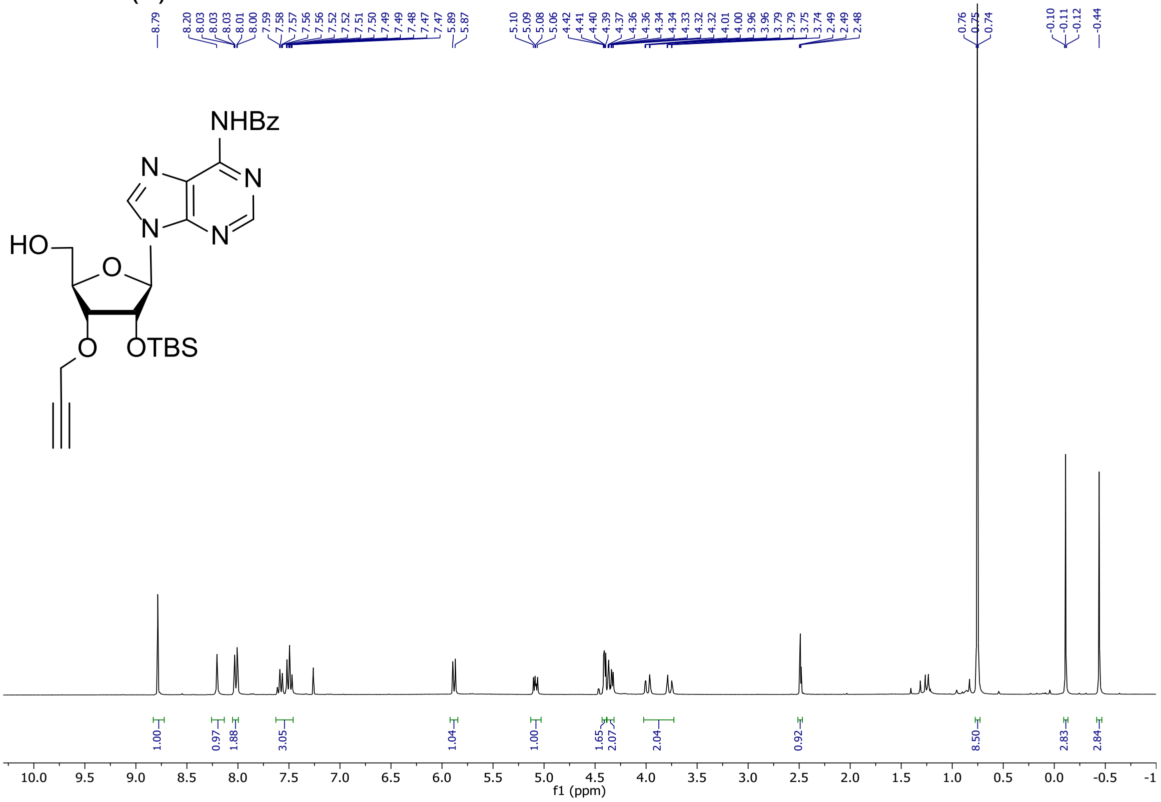


#### $^{13}C$ NMR (7)

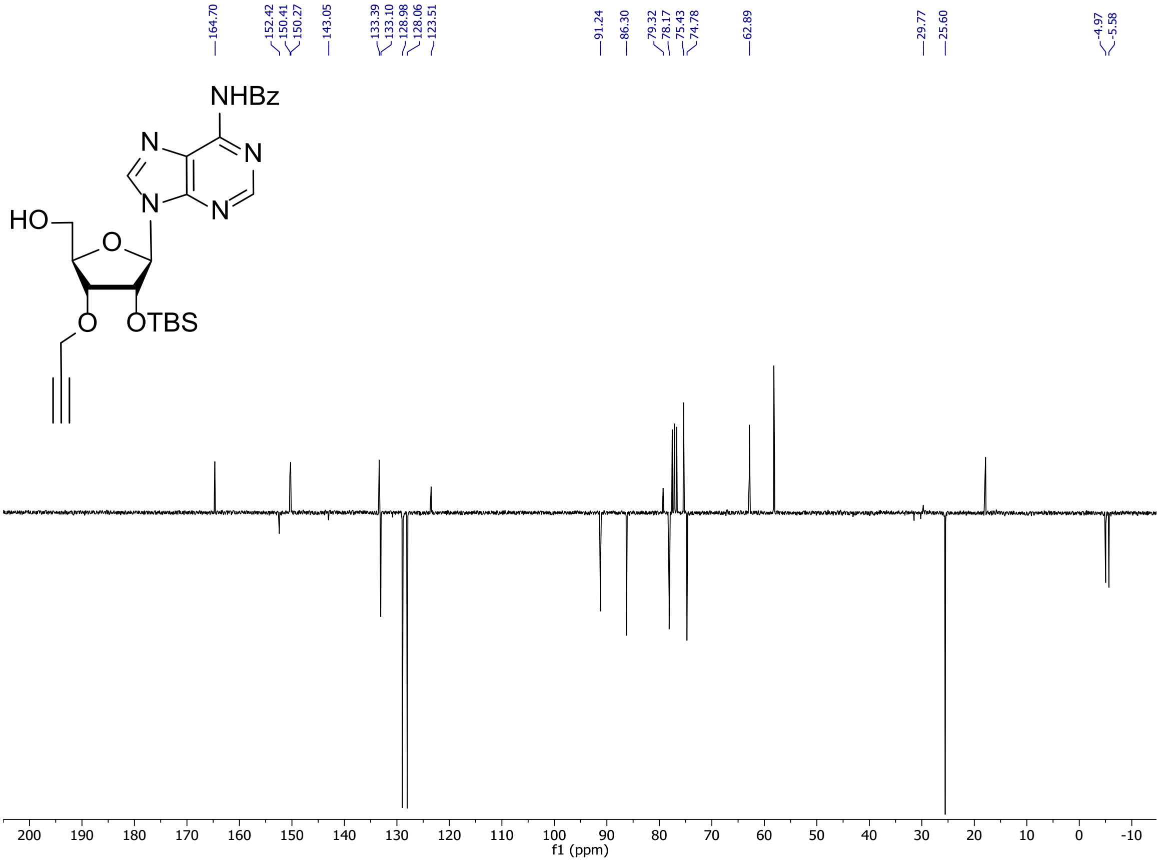

#### $^{1}$H NMR (**8**)

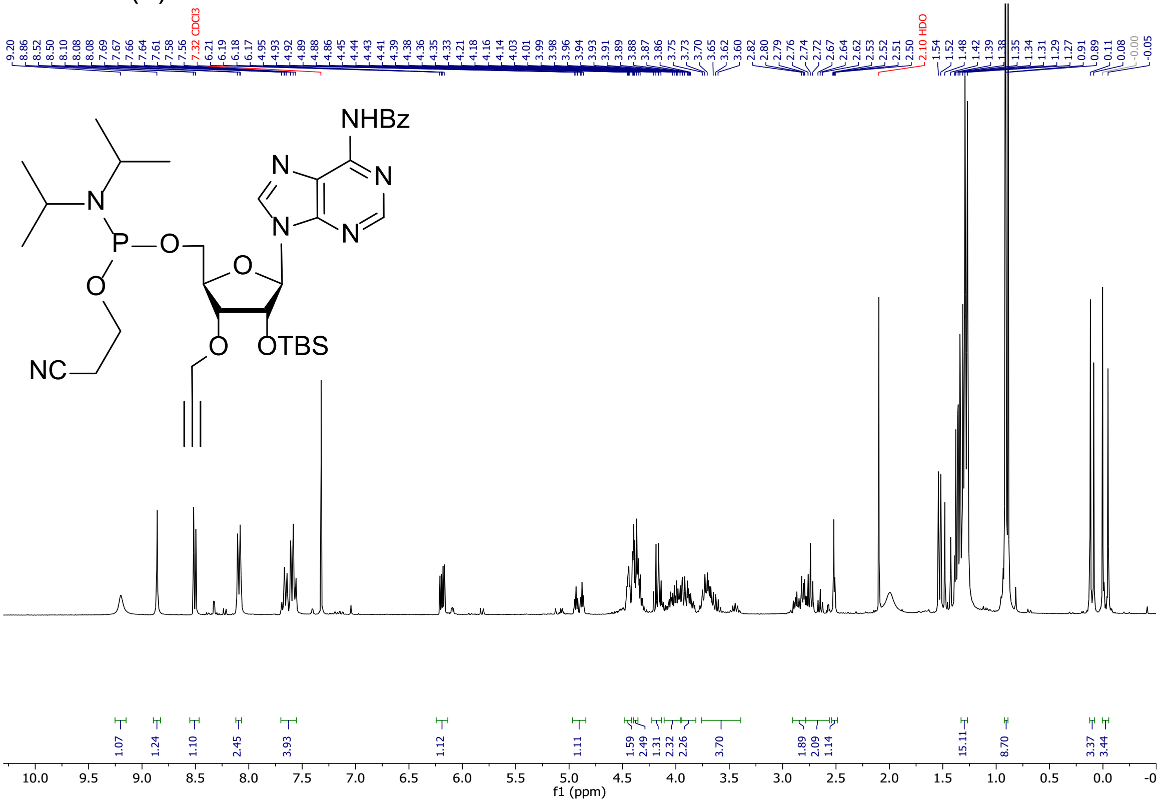

#### $^{13}$C NMR (**8**)

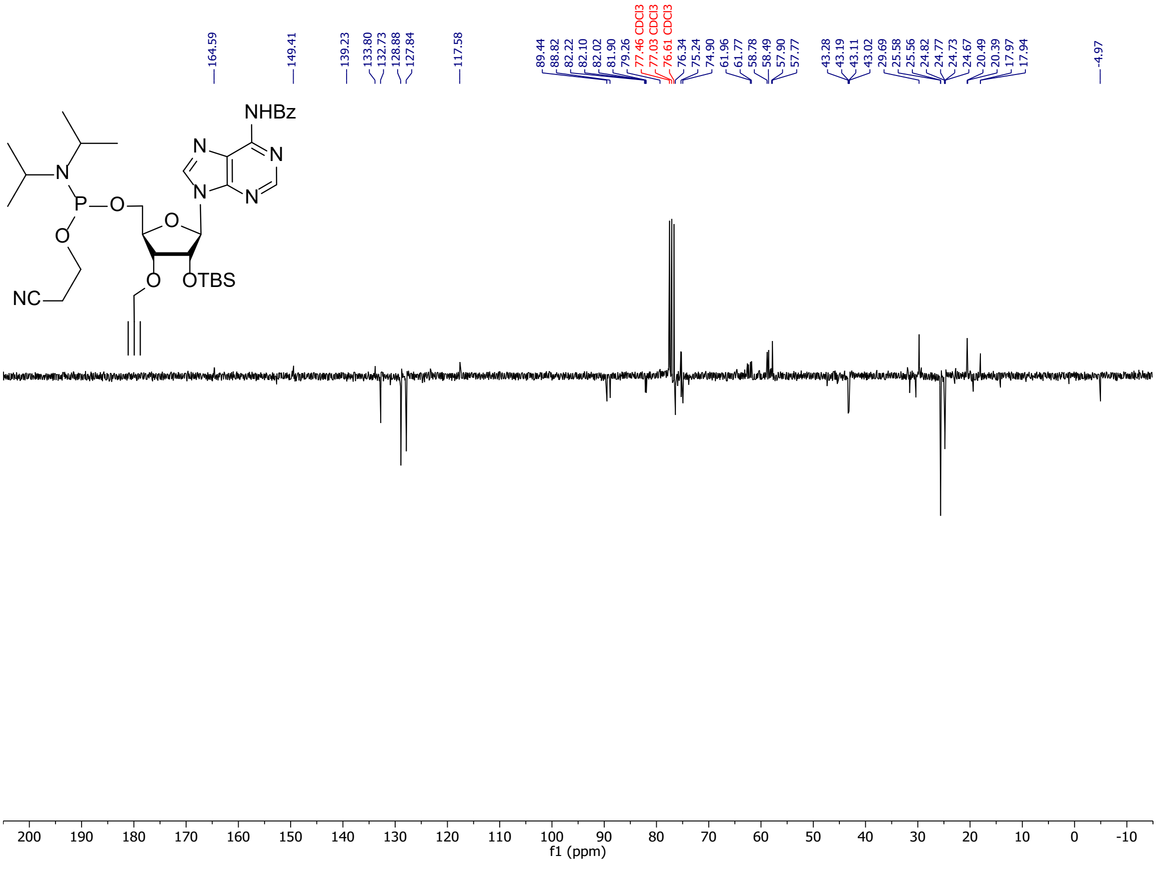

#### $^{31}$P NMR (**8**)

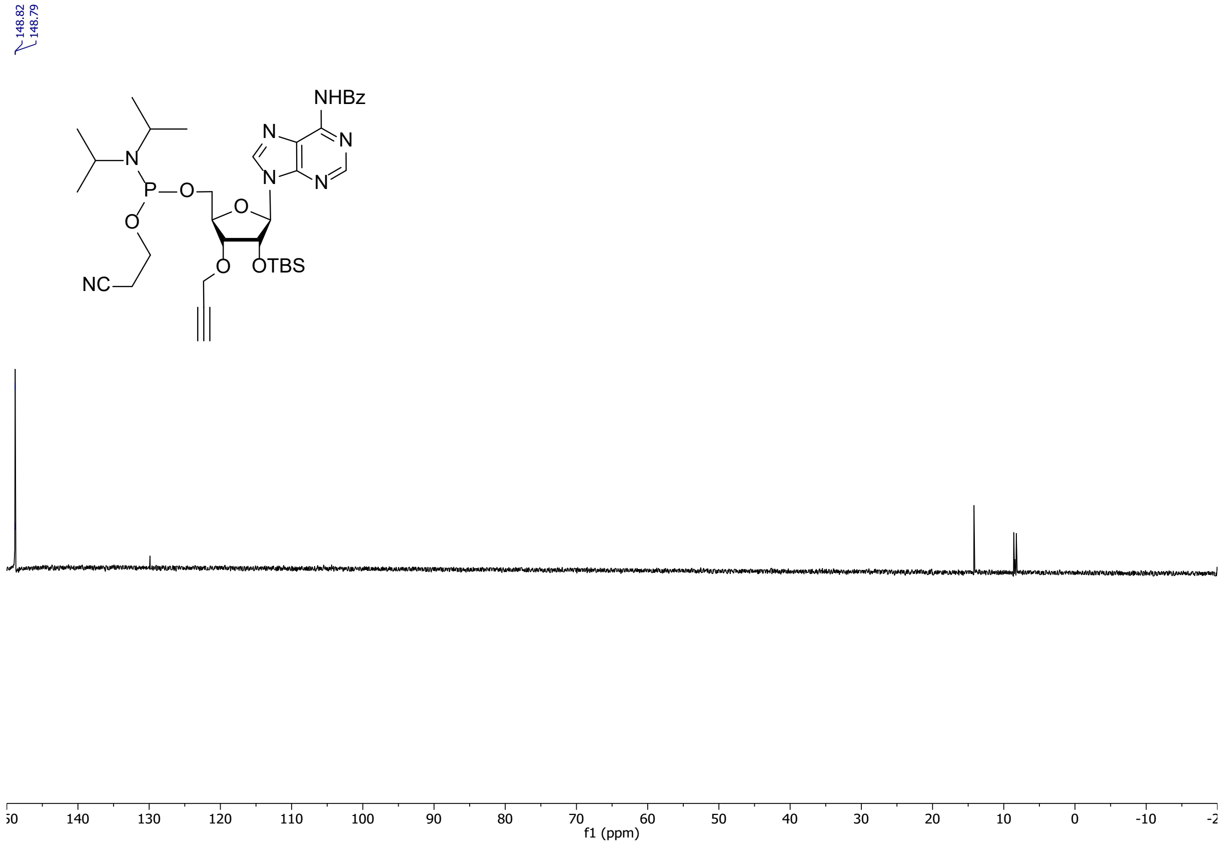


#### $^{1}$H NMR (**9**)

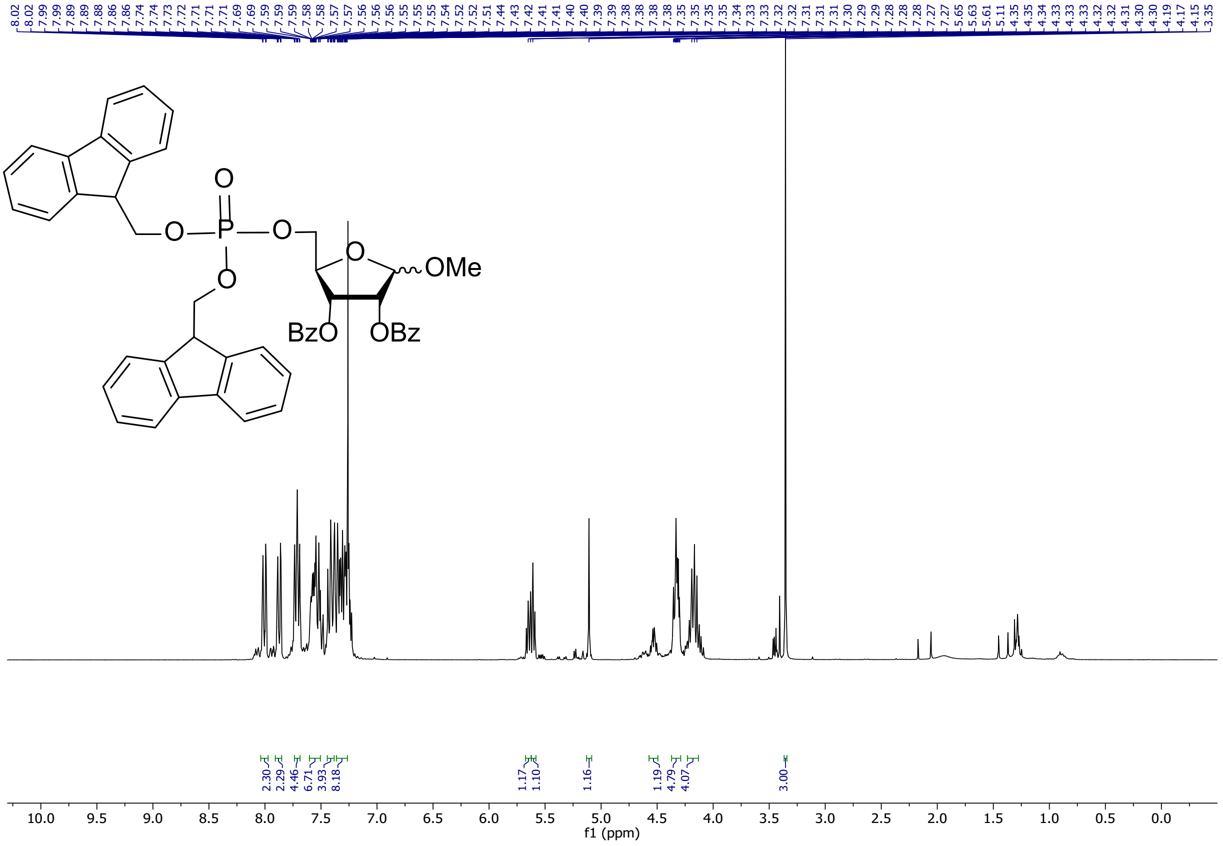

$^{13}C$ NMR (**9**)

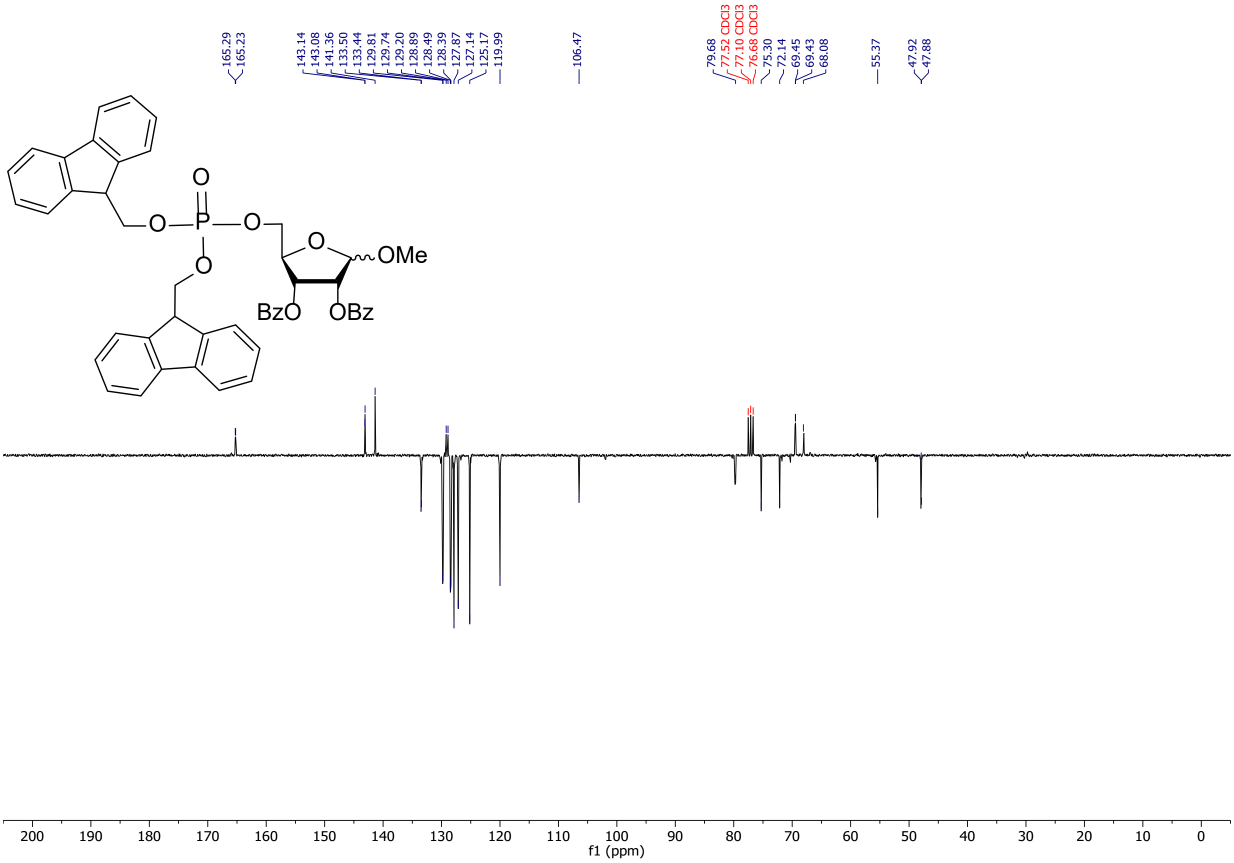


$^{31}P$ NMR (**9**)

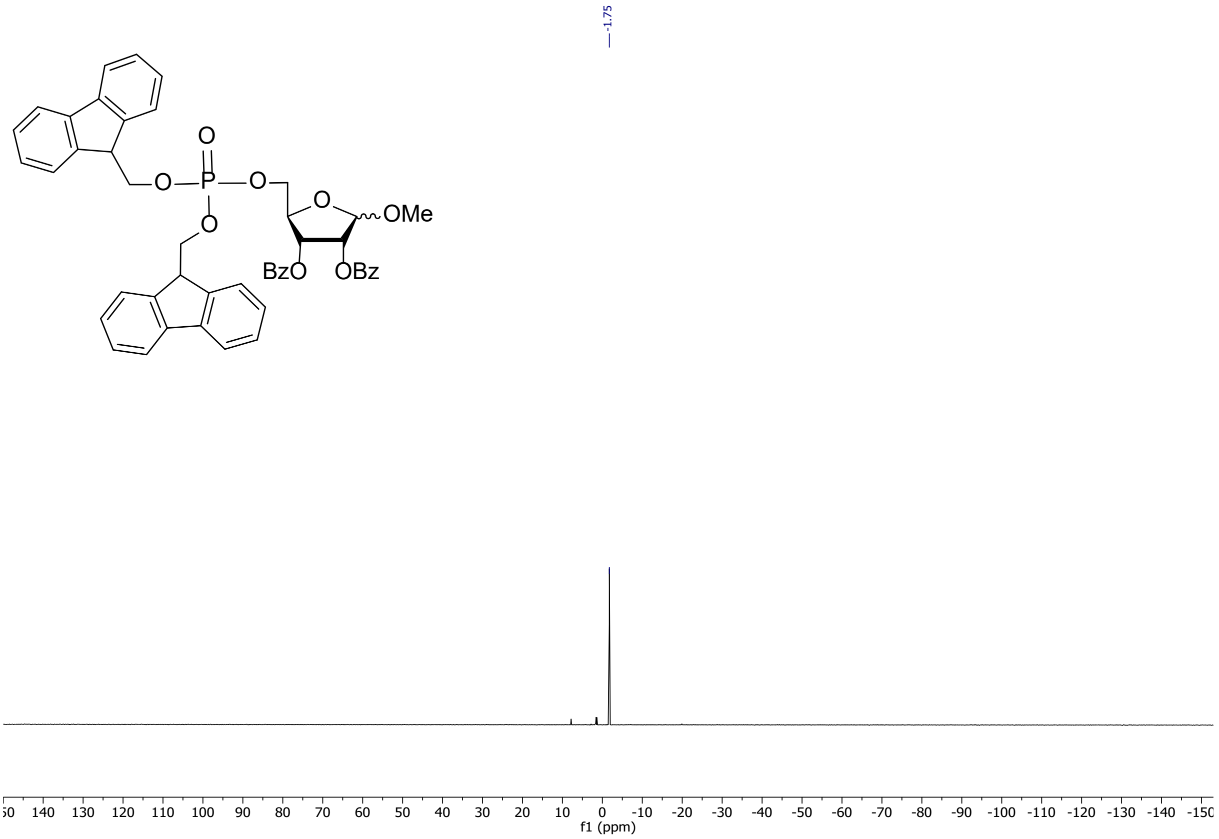

$^{1}$H NMR (**10**)

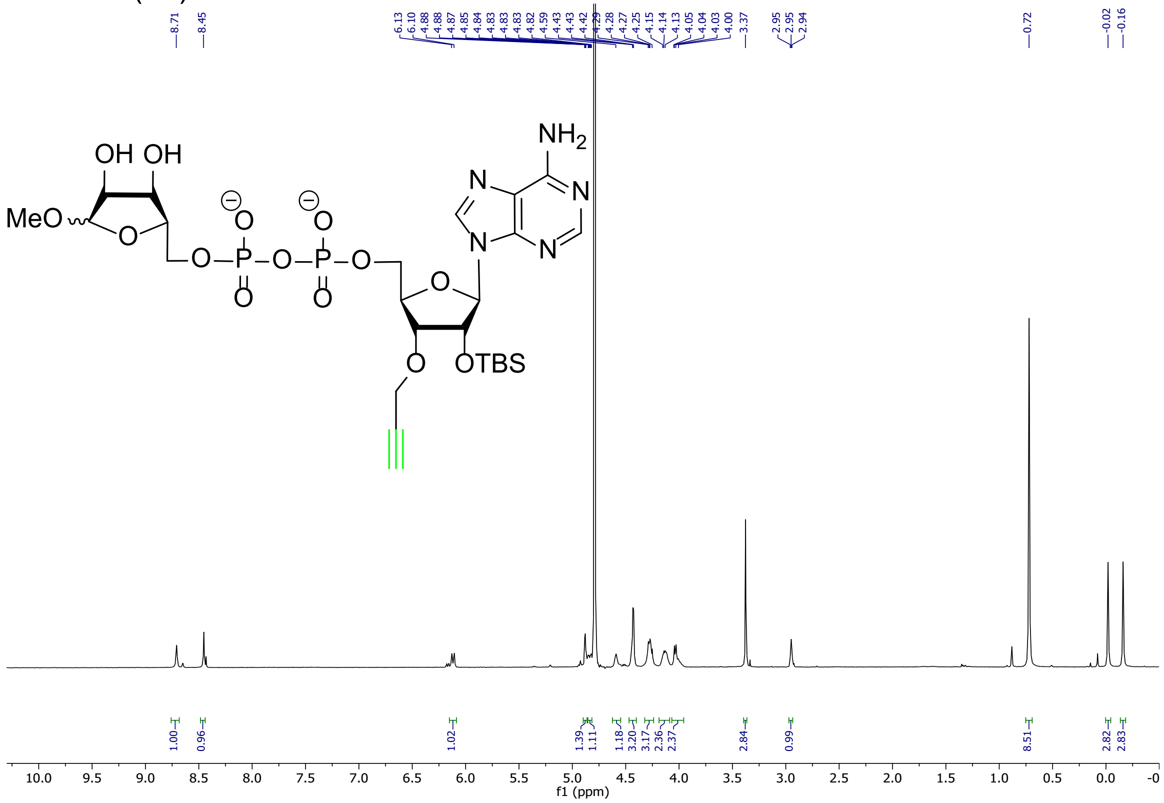

$^{31}$P NMR (**10**)

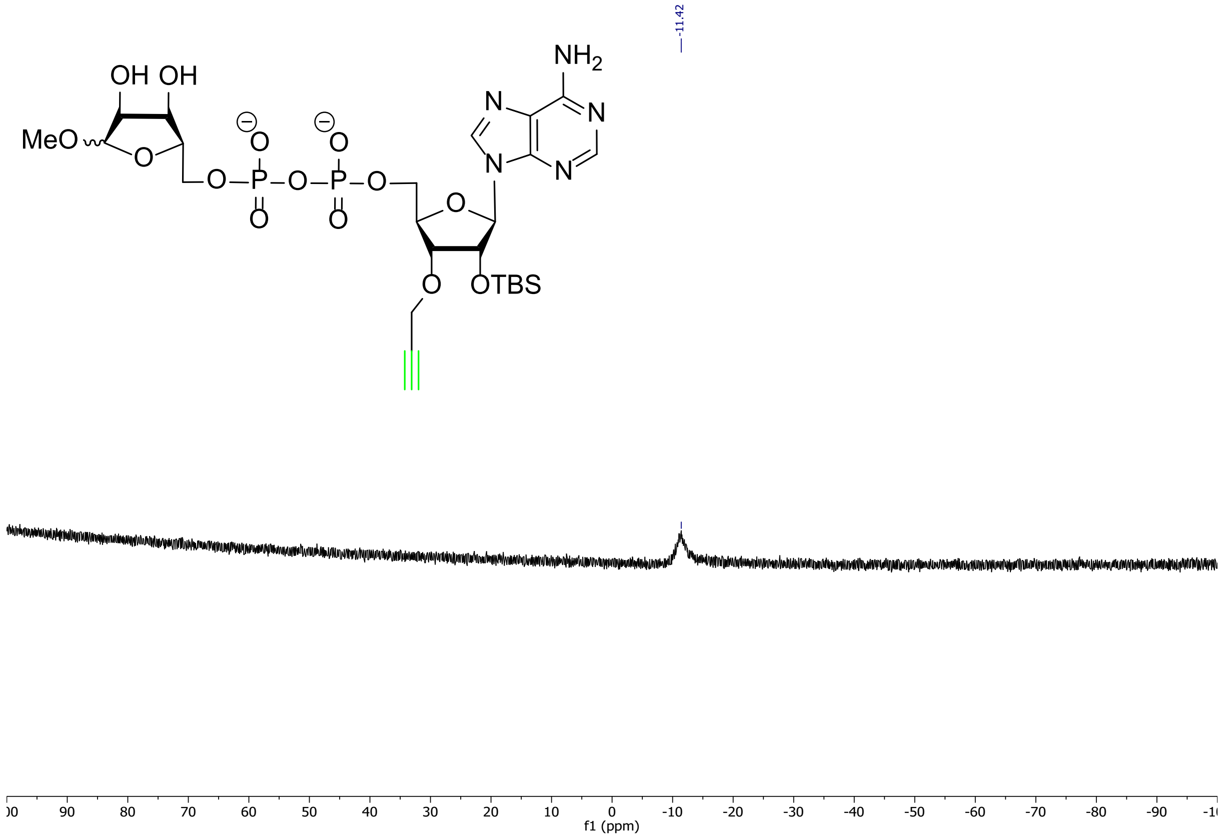
